# Olfactomedin4 marks luminal progenitor cells that give rise to secretory cell lineage in the mouse cervix

**DOI:** 10.64898/2026.04.28.721179

**Authors:** ShanmugaPriyaa Madhukaran, Yevgenia Fomina, Govindkumar Balagannavar, Elizabeth Payne, Jason Wilson, Lei Wang, Gary Hon, Maria Florian Rodriguez, Mala Mahendroo

## Abstract

Development of the female reproductive tract in mice occurs in early postnatal life. The current model identifies Trp63 as the master regulator that initiates differentiation of simple columnar Keratin 8+ epithelium in the cervix and vagina into a stratified squamous epithelium. Thereafter Trp63+ basal progenitors maintain cervicovaginal epithelial cell homeostasis and in the adult serve as the progenitor for hormone-regulated shifts in stratified squamous and secretory luminal cells. This model differs from the human in which two progenitors, one columnar and the other basal gives rise to secretory cells in the endocervix and stratified squamous epithelia in the ectocervix and vagina respectively. In the current study, we identify a population of Krt8+, Tp63- epithelial cells that are retained in the cervicovaginal epithelium during the postnatal developmental period and into adulthood. Single cell datasets from the cervices of adult mice, identify Olfactomedin 4 (Olfm4), as a unique marker of the Krt8+Trp63- population. Adult lineage tracing and reassessment of gene markers during postnatal development support a revised model in which two progenitors are delineated in the mouse cervix and vagina by PND15. Olfactomedin 4+ progenitors give rise to specialized secretory goblet cells, while Trp63+ basal progenitors give rise to stratified squamous luminal cells in the cervix and vagina of nonpregnant and pregnant mice. Consistent with the expansion of goblet cells in pregnancy, the Olfm4+ progenitor is highly proliferative in early pregnancy and progesterone regulates increased goblet cell differentiation. These findings reveal a previously unrecognized species similarity between mice and humans in which goblet cell and squamous keratinized cell subtypes are derived from two progenitor populations respectively.

**HIGHLIGHTS:**

- Two epithelial progenitors (Trp63 and Olfm4) populations are delineated in the cervix and vagina within the first two weeks of postnatal life.
- The Trp63+ progenitor gives rise to keratinized epithelial cells, whereas the Olfm4+ progenitor cells give rise to secretory goblet cells.
- lfm4 is not required for maintenance of the luminal progenitor or differentiation of goblet cells in the cervix and vagina during adulthood and pregnancy.
- In adults, progesterone promotes differentiation of Olfm4+ progenitors into goblet cells.

## INTRODUCTION

In the female reproductive tract, the cervical epithelium has a dual role; it facilitates sperm ascension necessary for fertilization and provides a protective mucosal barrier against ascending infections. During pregnancy, this barrier not only shields the maternal reproductive tract but also safeguards the fetus from infection and inflammation which can lead to preterm birth (PTB). In adults, cyclic hormonal changes drive the expansion of distinct cervical epithelial subtypes. One specialized subtype, the goblet cell, produces antimicrobials, protease inhibitors, and immunoglobulin transporters along with gel-forming mucins such as MUC5B and MUC5AC. In both mice and humans, reduced expression of MUC5B has been associated with increased rates of PTB (1, 2). In mice, goblet cells expand during pregnancy and have been well defined by canonical regulators of secretory differentiation such as SPDEF and CREB3L1 (3–6). Although mucin producing secretory cells are present in the human cervix, their identity as a canonical goblet cell has not yet been established. Given the conserved importance of mucin-mediated epithelial defense across other mucosal organ systems (7, 8) understanding the developmental origin and regulation of cervical mucin-producing cells is essential for defining how the cervical epithelium contributes to reproductive barrier function and pregnancy maintenance.

The cervical epithelium is anatomically divided into the endocervix, which is continuous with the uterine epithelium, and the ectocervix, which transitions into the vaginal epithelium. During early development in humans and mice, the cervical epithelium consists of a single-layered columnar epithelium marked by keratin 8 (KRT8). In humans, the endocervix remains columnar with glandular development by 16-21 weeks of gestation, whereas the ectocervix differentiates into a multilayered, stratified squamous epithelium under the control of the transcription factor, tumor protein 63 (TP63) (9, 10). In mice, tissue recombination experiments demonstrate that signals from the underlying mesenchyme induce the Krt8+ columnar epithelium lining the neonatal cervix to express Trp63 between postnatal days 1 and 5, driving epithelial stratification across both the endocervix and ectocervix (**Figure 1A**) (11–15). Additionally, in humans, the vaginal and ectocervical epithelium arise from the urogenital ridge epithelium (UGE) whereas the endocervix and upper female reproductive tract (FRT) arise from the Mullerian duct. In contrast, the entire mouse FRT, including the cervix and vagina, is derived from the Mullerian duct (16).

**Figure 1:**
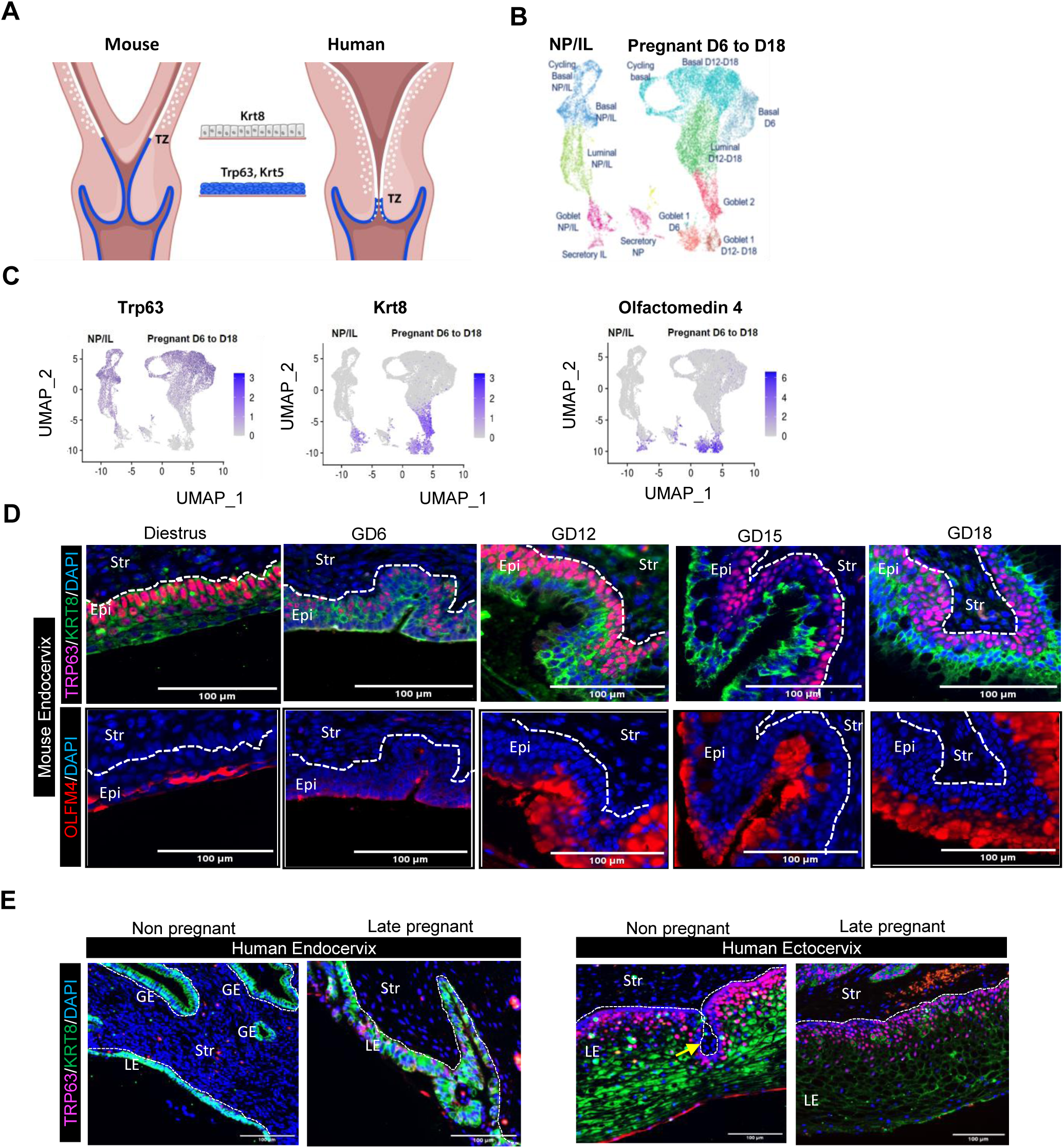
Expression of squamous and columnar epithelial populations in the mouse and human cervix across pregnant and nonpregnant states. A. Schematic comparison of the human and mouse female reproductive tract (FRT), highlighting the luminal and glandular columnar epithelial organization of the uterus, and the distinct columnar and squamous epithelial regions between the mouse and human cervix, with overlap of these subtypes localized at the transition zone (TZ). B. UMAP visualization of mouse cervical epithelium across reproductive states at non-pregnant (estrus and diestrus), pregnant (GD 6, 12, 15, 18) and in labor (IL) timepoints. Epithelial subtypes are color-coded to indicate basal, luminal (keratinocyte), secretory/goblet populations. (modified from (3)). C. Feature plots showing Trp63, Krt8 and Olfm4 (Olfactomedin 4) expression in mouse cervical epithelia. Expression intensity is displayed in purple. D. Immunofluorescence detection of TRP63 (red), KRT8 (green), and nuclei (blue) and OLFM4 (red) in mouse endocervical epithelium at non-pregnant (diestrus), pregnant (GD 6, 12, 15, 18) time points. Corresponding images of ectocervix and vaginal regions from the same animals are shown in Figure S1A. n≥3 biological replicates per time point. Scale bars =100 µm. E. Immunofluorescence detection of TRP63 (red), KRT8 (green) and nuclei (blue) in human non pregnant, and late-pregnant (34 weeks gestation) cervix. Images of endocervix and ectocervix regions are from the same individual; yellow arrow indicates stromal papillae. Luminal epithelial (LE) and glandular epithelia (GE) with n ≥ 3 biological replicates per group and scale bars = 100 µm.

Developmental distinctions in epithelial origin may explain why the adult human endocervix remains columnar, whereas the adult mouse endocervix is stratified squamous. These species-specific differences raise the question of how distinct progenitor populations, stratified squamous progenitors in the mouse endocervix and columnar progenitors in the human endocervix, can give rise to similar secretory epithelial subtypes with conserved functions. While epithelial cell-fate plasticity in the female reproductive tract was previously thought to be restricted to the postnatal period, accumulating evidence indicates that bipotent stem cells persist in the uterus and cervix into adulthood and in the cervix can generate both squamous and columnar epithelial lineages (17, 18). Maintenance of epithelial cell fate in the adult uterus is regulated in part by a balance in estrogen and retinoic acid signaling (19–21).

The focus of this study is to define the progenitor cell(s) that give rise to the cervical secretory goblet cell. Leveraging our single cell transcriptomic dataset from mouse cervical epithelium, we delineate markers that distinguish goblet cells from other epithelial subtypes (3). Among these, we identified olfactomedin 4 (Olfm4), a previously described marker of intestinal stem cells, as a distinguishing marker of cervical goblet cells (22). Using lineage tracing in reporter mice, we demonstrate the presence of a second progenitor population in the mouse cervix. This luminal progenitor is marked by Olfm4 by postnatal day 15 and gives rise to secretory goblet cells in the adult mouse cervix and vagina. We propose the Olfm4+ columnar progenitor is distinct from Trp63+ basal progenitors, which give rise to stratified luminal subtypes. Our findings reveal a novel progenitor lineage marker for goblet cells in the mouse cervix and vagina and highlight the epithelial diversity and plasticity of the adult lower female reproductive tract.

## RESULTS

### Identification of columnar and squamous markers in nonpregnant and pregnant mouse and human cervical epithelium

The adult mouse lower FRT epithelium consists primarily of a stratified squamous epithelium, in which a basal progenitor gives rise to cycle-dependent luminal cells (**Figure 1A**) (12, 16, 23, 24). Consistent with this model, our prior single-cell RNA sequencing studies identified luminal cell heterogeneity across nonpregnant, pregnant and laboring mice (**Figure 1B**) (3). The luminal populations were subclassified based on gene expression markers into i) secretory goblet cells, which synthesize gel-forming mucins (Muc5b, Muc5ac); and ii) a second secretory subtype predominant in nonpregnant (NP) cervix that expressed the transmembrane mucin (Muc1); and iii)keratinocyte-like cells (Dsg1a+, Dmkn+, Krt16+, Krt12+). Importantly, these results demonstrated that goblet cells are the primary luminal cells lining the mouse endocervix and ectocervix during pregnancy, labor and present in lower abundance during the diestrus phase of the reproductive cycle. Previous studies in pregnant individuals and mice suggest goblet cell dysfunction disrupts the mucosal barrier and increases risk of ascending infection-mediated PTB (2, 25). To better understand the lineage trajectories that give rise to this functionally important subtype, we characterized goblet cells using markers that distinguish them from other epithelial subtypes. As shown in the UMAPs in **Figure 1C**, goblet or secretory clusters lacked expression of the squamous marker *Trp63* and instead expressed the columnar cell marker, *Krt8*. Notably, olfactomedin 4 (*Olfm4*) was uniquely enriched in goblet cell clusters.

The spatial distribution of TRP63, KRT8 and OLFM4 proteins was assessed in the mouse endocervix, ectocervix and vagina across NP diestrus, pregnancy gestational days (GD) 6, 12, 15, and 18 (**Figures 1D and S1A).** TRP63+ cells were confined to the endocervical, ectocervical and vaginal basal layer adjacent to the stroma across all stages. KRT8+ cells were localized to luminal cell layers during diestrus and in pregnancy (**Figures 1D and S1A).** On GD6, a few TRP63+ KRT8+ double positive cells were present in the basal layer yet steadily declined in the later pregnancy timepoints. OLFM4 expression mirrored KRT8 expression in the luminal epithelial layers across all stages, consistent with the spatial localization and temporal pattern of goblet cells (**Figures 1D and S1A).**

We next performed a parallel analysis in the human endocervix and ectocervix from premenopausal or late pregnant individuals (**Figure 1E**). Both luminal and glandular epithelia of the endocervix were Krt8+ consistent with columnar cells. Interestingly, a few endocervical cells were also Trp63+ in the late pregnant cervix. The basal layers of the ectocervix were Trp63+ in both NP and late pregnant. The luminal layers were Krt8+ with some Trp63 Krt8 co-positive cells. In contrast to mice, OLFM4 protein expression was not detectable in the human endocervix or ectocervix (Figure **S1B**).

### OLFM4^+^ luminal cells undergo proliferation in early pregnancy

Olfm4 marks epithelial progenitor stem cells in the intestine (22, 26, 27) and prostate (28) where Olfm4 positive cells contribute to tissue regeneration and maintenance. In the mouse cervix, immunofluorescent co-staining of OLFM4 and the proliferation marker, KI67, identified proliferative OLFM4+ cells in luminal layers during secretory cell expansion on GD6 **(Figure S1C).** In contrast, in estrus and late pregnancy (GD15), proliferating cells were confined to the basal layer of both the endocervix and ectocervix, suggesting that basal cells are the predominant proliferative population at these stages. In early pregnancy (GD6), proliferation extended across all epithelial layers in both regions **(Figure S1C).** The presence of stage-specific proliferative OLFM4+ cells in the cervix, together with the established regenerative potential of OLFM4 marked cells in other tissues, led us to hypothesize that OLFM4 marks a luminal progenitor population in the cervical epithelium.

### Olfm4 expression is initiated during cervical postnatal development in mice

We sought to determine if Olfm4 is expressed during postnatal development of the FRT. To define the temporal pattern of OLFM4 expression during cervical development, we assessed its expression in combination with established markers, KRT8 (columnar) and TRP63 (squamous) at key postnatal timepoints in FRT development, PND 1, 5, 15, and 25 (**Figures 2A and S2).**

**Figure 2:**
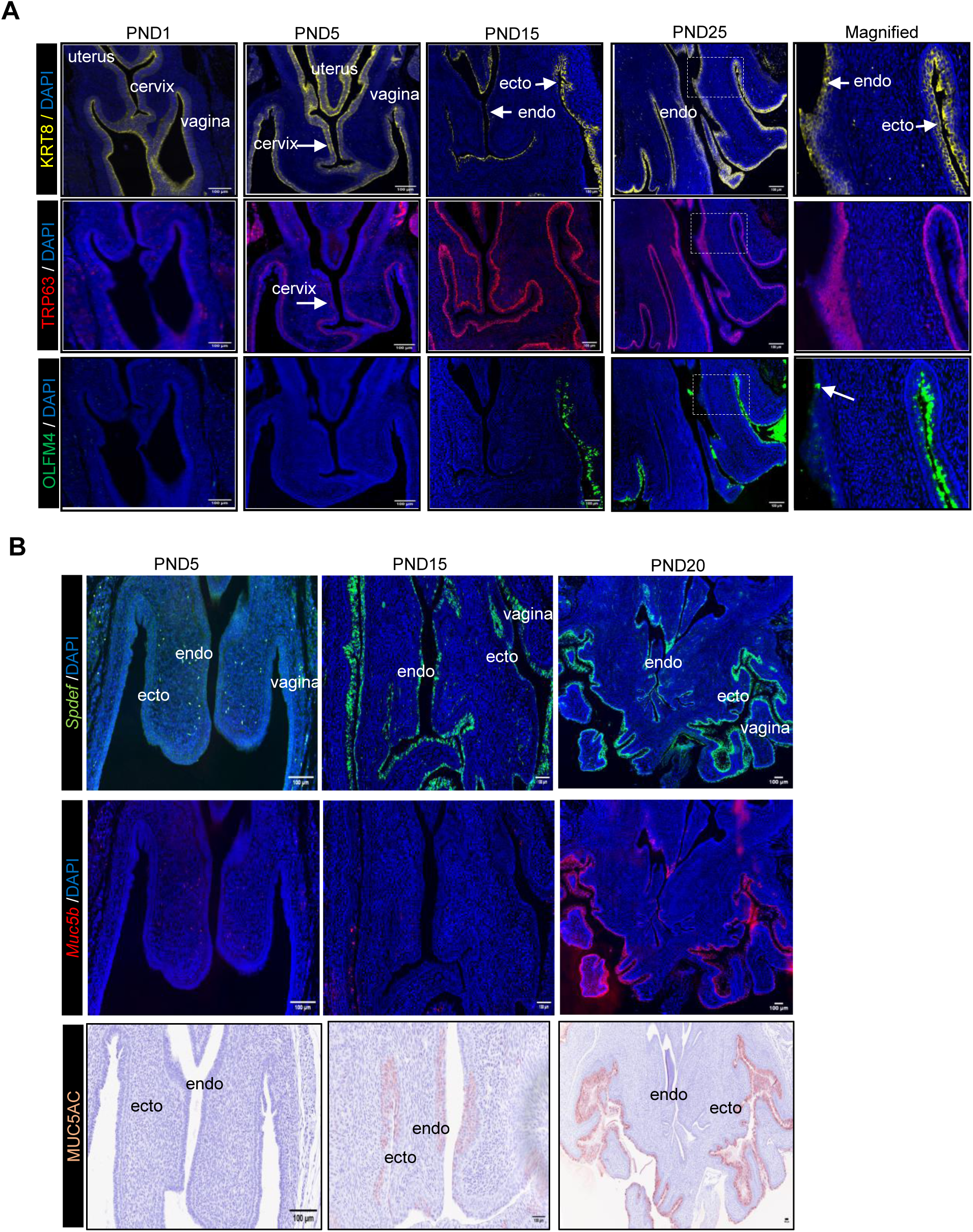
Olfm4 marks luminal cervical epithelia during postnatal FRT development. A. Immunofluorescence staining of the FRT for KRT8 (yellow), TRP63 (red), and OLFM4 (green) at postnatal days (PND) 1, 5, 15, and 25. The right-most panels shows a higher-magnification view of the PND25 cervix, highlighting spatial restriction of OLFM4 to the luminal columnar cells in the endocervix (white arrow), ectocervix and vagina; serial sections from the same animal were used for each antibody at each time point, with n=3 biological replicates per time point. Scale bars=100 µm. B. RNA expression of Spdef and Muc5b performed on the same tissue sections (signals displayed in separate panels for visualization) scale bar 100 µm. Immunostaining for MUC5AC protein in the FRT on postnatal days at PND5, PND15 and PND20 (n=2 biological replicates per time point). Scale bar=100 µm.

On PND1 the cervical epithelial layers are KRT8+ columnar epithelia. By PND5, all of the multilayered cervical epithelium was a KRT8+ epithelium and the basal layer of cells co-expressed TRP63. OLFM4 expression was absent at both PND1 and PND5, thus does not appear to be an early marker of cervical epithelial differentiation. Between PND15 and PND25, TRP63 expression was predominant in the basal layers of epithelial cells, while a thin luminal layer of cells remained KRT8+. Among the KRT8+ luminal cells, a subset expressed OLFM4 in the endocervix with a greater number of KRT8+OLFM4+ cells in the ectocervix and vaginal epithelia (**Figures 2A, S2B).** The KRT8+OLFM4+ cells also expressed goblet cell markers including *Spdef,* (SAM pointed domain containing ETS factor), a transcription factor required for goblet cell differentiation (3) and the secretory mucins, *Muc5b* and MUC5AC (**Figure 2B**) suggesting cell-fate specification of goblet progenitors is established by PND15. These findings support a stepwise delineation of two distinct epithelial progenitor populations in the postnatal period with basal-localized TRP63+ and a luminal-localized KRT8+/OLFM4+ population. In contrast to the cervix, the expression of Olfm4 and goblet cell markers are not detected in the uterine epithelium in the postnatal period (**Figure S2C**). This observation is consistent with published scRNA-seq datasets of the uterus between postnatal days 1-15 in which < 2.6 % of cells had detectable cell counts for Olfm4 or Spdef **Figure S2D** and (29).

### Krt8+ progenitors give rise to goblet cells in the cervix and vagina

Given the observation that the luminal layers of cervical epithelia do not gain Trp63 expression during the FRT developmental period and express Krt8, we tested the potential of a Krt8+ progenitor cell to give rise to secretory goblet cells in the adult mouse cervix before and during pregnancy. Lineage tracing studies were conducted in the Krt8-Cre^ERT2^; Rosa26^tdTomato^ females as described in (**Figure 3A**). In the cervix and vagina, tdTomato+ cells were primarily confined to luminal layers. The tdTomato+ cells expressed secretory goblet cells markers, *Spdef* and *Muc5b* during diestrus and pregnancy timepoints confirming that Krt8+ progenitor cells give rise to goblet cells (**Figures 3B and S3).** These findings provide support for the presence of a cervical columnar Krt8+ progenitor as the precursor for secretory goblet cells.

**Figure 3:**
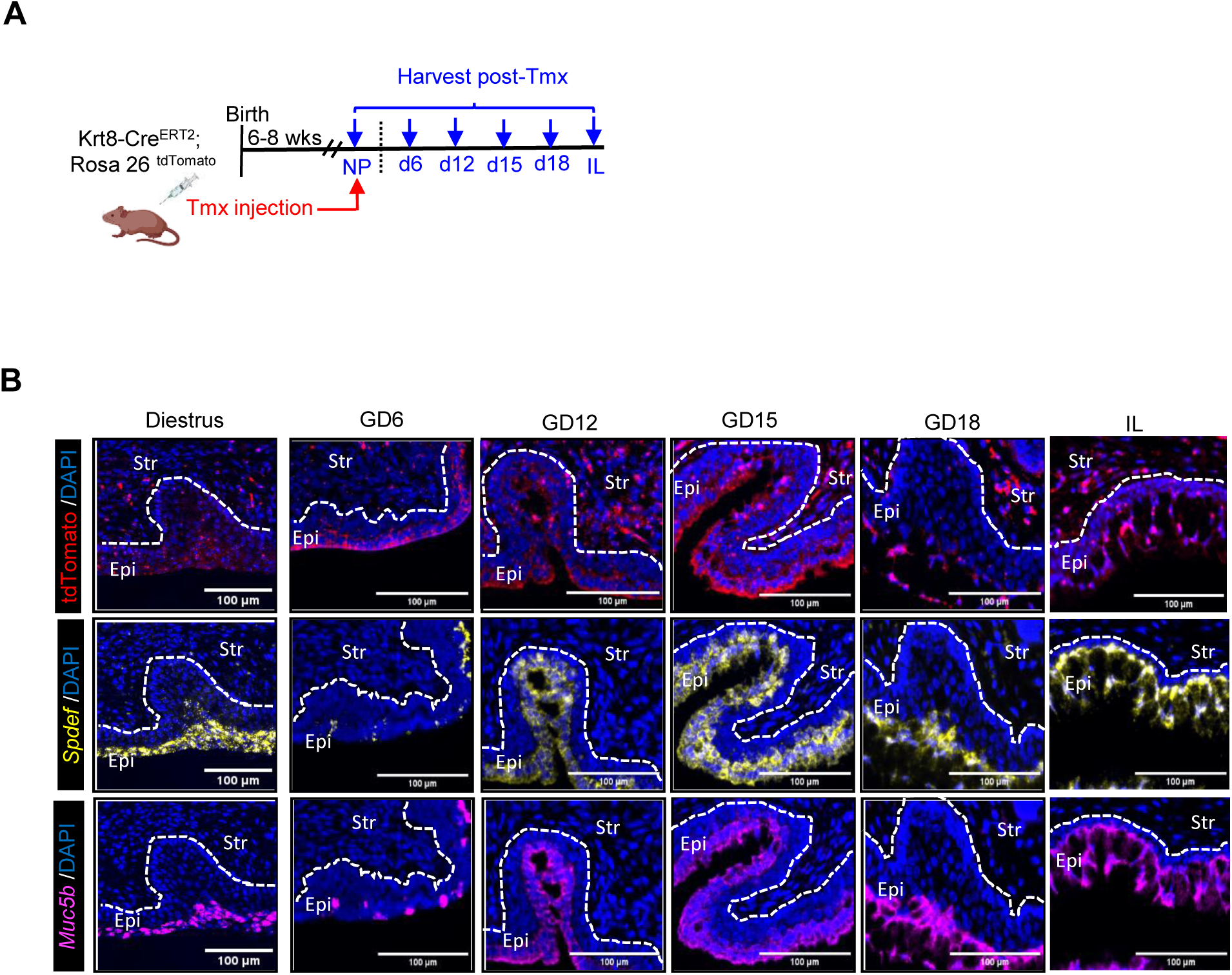
Krt8⁺ progenitor cells give rise to goblet cells in the cervix. A. Schematic of lineage-tracing strategy using Krt8-Cre^ERT2^; Rosa26^tdTomato^ mice with tissues collected across nonpregnant and early-to late-pregnancy: gestation day (GD) 6, 12, 15, 18 and in labor (IL). B. RNA expression in the endocervix for tdTomato (red), Spdef (yellow), and Muc5b (pink) across diestrus, pregnancy stages and in labor. Spdef and Muc5b were co-stained in the same section, with each marker displayed in separate panels, whereas tdTomato was detected on a separate section. Epi-epithelial and Str- stromal regions are indicated. Images are representative of n=1 biological replicate per time point and scale bars =100 µm.

### Olfm4+ luminal progenitors represent a self-renewing epithelial population that give rise to cervical secretory cells

To evaluate the progenitor potential of Olfm4-expressing cells in the cervix, we conducted lineage tracing studies using Olfm4-IRES-eGFPCre^ERT2^;Rosa26^tdTomato^ reporter mice (22). Nonpregnant Olfm4Cre^ERT2^; R26^tdTomato^ mice received a single intraperitoneal injection of Tamoxifen (Tmx;1mg/g body weight) (**Figure 4A**). Following tamoxifen induction, mice were either maintained as nonpregnant or timed-mated, and the cervical tissues were collected for spatial assessment of tdTomato expression and lineage-specific mRNA or protein markers. At all-time points in pregnancy and nonpregnancy, tdTomato+ cells were restricted to the apical luminal layer of the endocervix (**Figure 4B**), ectocervix and vagina **(Figures S4A and B)**, indicating that these cells do not arise from a basal progenitor population but instead originate from an Olfm4+ progenitor compartment.

**Figure 4.**
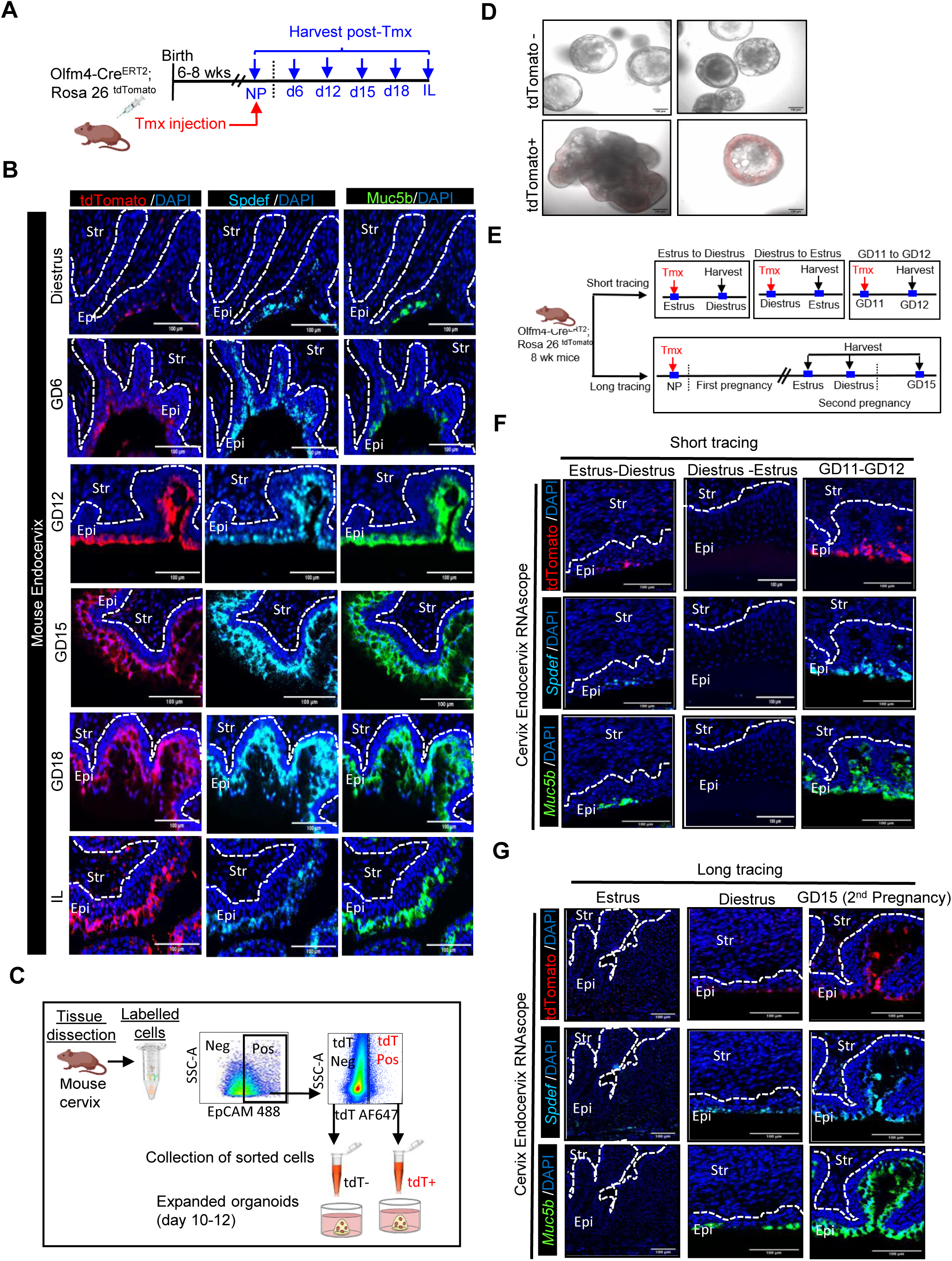
OLFM4+ progenitors represent a self-renewing epithelial population that give rise to cervical secretory cells. A. Schematic of lineage tracing strategy using Olfm4-Cre^ERT2^; Rosa26^tdTomato^ mice. B. RNA expression of tdTomato+ (red), Spdef (cyan) and Muc5b (green) in endocervical tissue from nonpregnant (diestrus) and pregnancy (GD 6,12,15, 18 and IL). All three markers were detected in the same section, with each signal displayed in separate panels. Images are representative of n≥ 3 biological replicates per time point and scale bars =100 µm. C. Schematic of workflow used to generate organoid cultures from tdTomato negative and tdTomato positive epithelial populations from the cervix of nonpregnant diestrus Olfm4-Cre^ERT2^; Rosa26^tdTomato^ mice. D. Organoids generated from flow cytometry sorted tdTomato+ and tdTomato- epithelial populations. tdTomato+ derived organoids retained tdTomato signal, whereas tdTomato-progenitors formed organoids that did not express tdTomato. n = 3 biological replicates and scale bars =100 µm. E. Schematic of short-term and long-term lineage-tracing strategy using Olfm4 Cre^ERT2^; Rosa26^tdTomato^ mice (8-week-old females). Short-term tracing (top): Tamoxifen (Tmx) was administered at estrus, diestrus, or GD11, and tissues were collected 24-48 hrs later at the subsequent time points (estrus to diestrus, diestrus to estrus, GD11 to GD12). Long-term tracing (bottom): Tmx was administered at the NP stage, followed by a first pregnancy. After the first pregnancy, tissues were collected at two stages of the estrous cycle, or on GD15 of a second pregnancy. F. RNAscope images of endocervix from short-term tracing experiments. Panels indicate tdTomato⁺ (red), Spdef (cyan), Muc5b (green) and nuclei (blue). In short-term tracing, tdTomato⁺ cells are present and co-localize with goblet markers during diestrus (first column) and on GD12 (third column). All three markers were stained in the same section and are displayed in separate panels. Representative images from n = 2 biological replicates per condition. Scale bars = 100 µm. G. RNAscope images of endocervix from long-term tracing experiments. Panels indicate tdTomato⁺ (red), Spdef (cyan), Muc5b (green) and nuclei (blue). In long-term tracing, tdTomato⁺ cells persist and give rise to secretory goblet cells that express Spdef and Muc5b in diestrus (second column) and in a second pregnancy at GD15 (third column). All three markers were stained in the same section and are displayed in separate panels. Representative images from n = 2 biological replicates per condition. Scale bars = 100 µm.

A gradual increase in luminally localized tdTomato+ cells was observed from GD6 to GD18 and in labor (**Figure 4B**). The spatial distribution of tdTomato+ expression in the cervix closely mirrored endogenous OLFM4 expression (**Figure 1D**), confirming reporter fidelity. RNAscope analysis also demonstrated overlap of cervical tdTomato with *Spdef* (6) as well as the secretory mucin *Muc5*b, providing further support that Olfm4+ progenitor cells give rise to the secretory goblet cells (**Figure 4B**). As an orthogonal approach, organoid cultures were derived from flow-sorted tdTomato^+^ and tdTomato^-^ cervical epithelial cells (**Figure 4C**). Organoids derived from tdTomato+ cells consistently retained tdTomato fluorescence and expression over multiple passages, whereas organoids derived from tdTomato- cells remained negative (**Figures 4D and S4C).** Despite the demonstration that organoids are derived from progenitor cells with lineage specificity, organoids from both tdTomato positive or tdTomato negative progenitors expressed Trp63+ cells and Krt8+ cells. This suggests cellular plasticity under the organoid culture conditions and warrants future investigation. Organoid growth efficiency was reduced in the tdTomato positive as compared to tdTomato negative organoids **(S4C).** Together, the lineage tracing data identify an Olfm4+ luminal progenitor population in the cervix that undergoes marked expansion early in pregnancy and subsequently differentiates into goblet cells throughout gestation.

Our previously reported single cell transcriptomic dataset from NP mice was generated from pooled estrus and diestrus samples (3). Within this dataset, two secretory epithelial populations were identified. One population was classified as canonical goblet cells, while the second was designated a non-goblet secretory population based on low Spdef expression and absence of Muc5b expression (3). To distinguish between two secretory cell types or transition states, Olfm4Cre^ERT2^; R26^tdTomato^ mice were assessed at proliferative (estrus), early secretory (metestrus) and secretory (diestrus) stages of the nonpregnant cycle for the presence of tdTomato+ cells and for *Spdef, Muc5b* and *Muc1* expression in the endocervix (**Figure S4D**). Consistent with their progenitor potential, tdTomato+ cells were present at all three stages of the cycle, with the highest abundance observed in diestrus. Integrated analysis of secretory marker expression across phases suggests a stepwise differentiation of Olfm4+ progenitors from an early/intermediate in estrus/metestrus to fully differentiated goblet cells in diestrus. This dynamic differentiation program positions Olfm4+ luminal progenitors as a shared unipotent progenitor pool for cervical goblet cells during both non-pregnancy and pregnancy.

In addition to the above-described regime for activation of the Cre reporter, tdTomato+ cells that express goblet cell markers were observed after short term (24-28hrs) and with long term (2nd pregnancy) tracing in which tdTomato+ cells continued to give rise to goblet cells in a second pregnancy (**Figures 4E-G**). Finally, tdTomato+ cells persisted after ovariectomy and expanded with progesterone but not estrogen treatment (**Figure S4E**). Collectively, these findings support the conclusion that Olfm4+ luminal progenitors represent a self-renewing epithelial population with a sustained capacity to generate cervical goblet cells across the reproductive cycle.

### Olfm4 is not required for goblet cell development

In addition to marking Lgr5+ intestinal stem cells, Olfm4 is a glycoprotein produced by epithelia and neutrophils with conserved roles in mucosal immunity across tissues, including the gastrointestinal and respiratory tracts (30–35). Since Olfm4 expression is initiated postnatally and up-regulated in the mouse cervix during pregnancy (**Figure 1D**), we asked whether Olfm4 is also required for cervical goblet cell differentiation and mucosal barrier function. Using mice with a targeted Olfm4 deletion (35) (), histologic analyses of nonpregnant and pregnant mice (GD 6-18) revealed morphologically similar secretory cells in the endocervix (**Figures 5A and S5A)**. Consistent with this, expression of goblet cell transcription factor Spdef and mucin Muc5b were comparable between genotypes (**Figures 5B and S5B),** indicating that Olfm4 is not required for cervical goblet cell differentiation. To assess barrier function in Olfm4 deficient mice, we introduced an ascending vaginal infection with pathogenic *E. coli* (serotype O55; ATCC) at GD15. Preterm birth rates were not significantly different between WT and Olfm4-deficient mice (p=0.198, Fisher’s exact test) **(Table S1)**. Next, we used bioluminescent *E. coli* to track bacterial ascension (34), with WT and Olfm4-deficient mice showing similar vaginal colonization at 0 hours and equivalent spread of *E.coli* to the uterus at 12 hours post-inoculation (**Figure 5C**). Inflammatory responses were further evaluated by intraperitoneal LPS administration (5mg/kg) at GD 15. No significant differences were observed in the expression of proinflammatory genes in the cervix (top panel) or uterus (bottom panel) **(Figure S5C)**.

**Figure 5:**
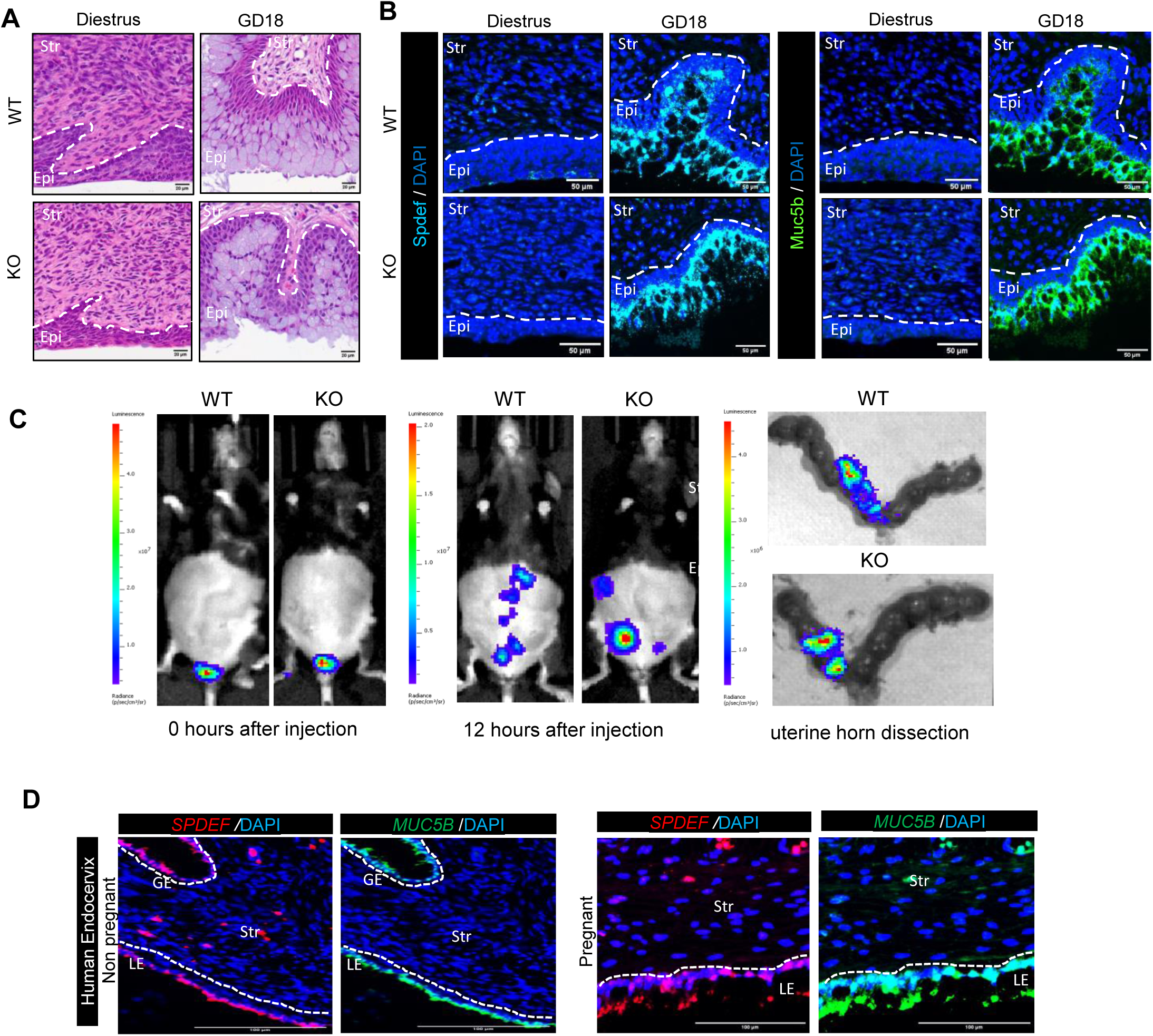
OLFM4 expression is not required for cervical goblet cell differentiation or function. A. Hematoxylin and eosin (H&E) staining of the endocervical canal in Olfm4 wild-type (WT) and knockout (KO) mice during diestrus and at gestational day 18 (GD18). Images are representative of n= 2 biological replicates per genotype and time point. Scale bars =20 µm. B. RNAscope staining for transcription factor Spdef (cyan) and Muc5b (green) in WT and KO endocervical tissues at nonpregnant diestrus and gestational day 18 (GD18). Both markers were detected in the same section, with each signal displayed in separate panels. Images are representative of n= 1 biological replicates per genotype and time point. Scale bars =50 µm. C. Inoculation with bioluminescent, non-pathogenic *E. coli* (10LJ CFU) at 0 and 12 hours, followed by dissection of the uterine horns at 12 hours post-inoculation in the WT and KO mice. Images are representative of n=3 biological replicates per time point. D. RNA expression of SPDEF (red), MUC5B (green), DAPI (blue) in human endocervix during nonpregnant and late pregnant states. Both markers were co-stained on the same section and are shown in separate panels. n=3 biological replicates per timepoint. Scale bars =100 µm.

Overall, these findings indicate that Olfm4 deficiency does not significantly affect cervical goblet cell differentiation, susceptibility to ascending *E. coli* infection, rates of PTB, or proinflammatory responses to systemic LPS exposure. Consistent with this observation in mice, goblet cells in the human endocervix express canonical markers Spdef and Muc5b (36) (**Figure 5D**) but do not express Olfm4 (**Figure S1B**). Thus, while Olfm4 marked progenitors give rise to goblet cells, Olfm4 itself is dispensable for goblet cell differentiation and maintenance of the cervical mucosal barrier. The absence of a strong phenotype in Olfm4-deficient mice likely reflects redundancy with other secreted factors, as has been reported in the intestine and other mucosal tissues.

### Progesterone induces Olfm4+ progenitor differentiation into goblet cells

Progesterone is an established driver of secretory cell expansion and mucin synthesis while estrogen promotes keratinocyte differentiation in the lower FRT (37–42). We thus sought to evaluate the impact of progesterone on the proliferation and differentiation of the Olfm4+ luminal progenitor. To test this, female NP mice were ovariectomized (OVX) to deplete endogenous ovarian hormones and subsequently treated for 15 days with vehicle, estradiol (E2), or progesterone (P4). Cervical epithelial populations were then evaluated to assess the effects of each hormone (**Figure 6A**).

**Figure 6:**
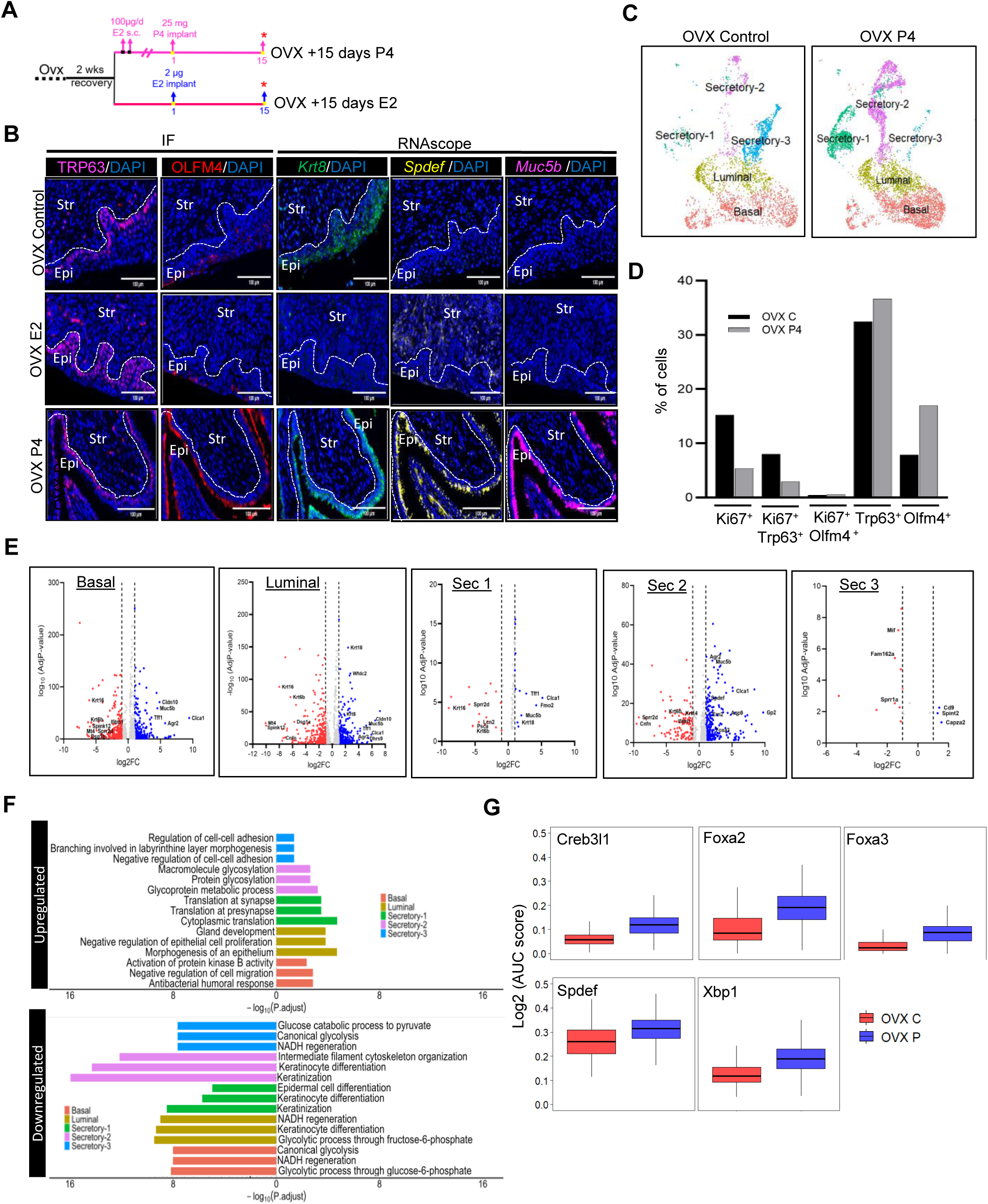
Progesterone induces Olfm4_⁺_ progenitor differentiation into goblet cells. A. Schematic of hormonal treatment in ovariectomized (OVX) mice. B. Immunofluorescence staining for TRP63 (basal; pink), OLFM4 (red), nuclei (blue). RNAscope staining for *Krt8* (green), *Spdef* (yellow) and *Muc5b* (pink). TRP63 and OLFM4 were detected on serial sections, whereas *Krt8*, *Spdef* and *Muc5b* were co-stained on the same section and displayed in separate panels. C. UMAP visualization of single-cell cervical epithelial populations from OVX Control and OVX P4 treated mice, with n=1 library per group and each library is comprised of a pool of 9 cervices. D. Percentage of proliferative epithelial cell subpopulations in OVX (black) and OVX P4 (gray) cervices. Bar graph shows the percentage of proliferating (Ki67^+^) and total Trp63^+^ basal cells and Olfm4^+^luminal cells. As data was obtained from a single library per group, statistical analysis was not performed. E. Volcano plots of differentially expressed genes (DEG) in OVX P4 versus OVX control within each epithelial subtype (basal, luminal, secretory -1, -2, -3) with selected upregulated and downregulated genes labeled. F. Gene ontology enrichment analysis of DEGs within each epithelial cluster indicating Downregulated and upregulated pathways in the OVX+P4 group are indicated. G. SCENIC analysis of predicted transcription factor regulons induced in the Sec-2 epithelial cluster of Ovx P4 samples.

Spatially distinct epithelial subpopulations were noted with increased OLFM4 protein expression with P4 treatment and increased expression of keratinocyte markers, *Dsg1a+, Krt16+* with E2 treatment (**Figures 6B and S6A).** TRP63 expression was consistently maintained across all treatment groups and predominantly localized to the basal layers. OLFM4+ cells were confined to the luminal layers with a robust increase in OLFM4+ cells in the P4 treated but not E2 treated mice. In general, cells expressing *Krt8* transcripts overlapped with OLFM4+ luminal epithelium in the OVX control and both treatment groups. *Krt8* expression was lost under E2 treatment but induced by P4, with robust expression across multiple luminal layers. RNAscope staining for goblet cell markers, *Spdef* and *Muc5b* demonstrate the OLFM4+ cells induced by P4 treatment express these markers (**Figure 6B**).

We next sought to determine, at a single-cell resolution, the P4-driven transcriptomic changes in epithelial cell states that lead to goblet cell differentiation. Using the 10X Genomics platform, scRNA-seq libraries were generated from OVX control (OVX C) and OVX P4 treated cervices. After quality control filtering, canonical markers for epithelial, immune, and stromal cells were utilized to annotate the dataset. In the current analysis, non-epithelial cells were computationally excluded from downstream analysis. The remaining epithelial cells were clustered into distinct epithelial subtypes using Seurat-predicted markers. Five epithelial clusters were identified in both OVX C and OVX P4 treated conditions: basal, luminal, secretory 1, secretory 2, and secretory 3 (Sec 1, Sec 2, Sec 3) (**Figure 6C**). As shown in **Table S2**, the OVX P4 treated group had a marked increase in Sec 1 (20 vs 4%) and Sec 2 (23 vs 9%) clusters, a reduction in Sec 3 (1 vs 24%) cluster, and only modest changes in the percentage of cells in the basal (37.5 vs 41%) and luminal clusters (18 vs 22%) relative to the control group.

A comparison of proliferative *Ki67+* populations between control and P4-treated groups indicated an overall decline in the percent of proliferative cells with P4. While there was no change in the percentage of proliferating Olfm4 (*Ki67+Olfm4+)* cells, there was a marked reduction in the percentage of proliferating *Trp63 (Ki67+Trp63+)* cells in the P4-treated group. Thus, P4 does not induce proliferation of the Olfm4+ progenitor but does limit proliferation of the basal Trp63+ progenitor (**Figure 6D**). The percentage of total *Olfm4+* cells increased with P4 treatment relative to OVX C, consistent with the observed increase in goblet cells as seen in **Figure 6B**. Gene markers that distinguish basal/luminal cells were present in both control and P4-treated groups as seen in the UMAP visualization **(Figure S6B).** Sec clusters 1 and 2 which are enriched in the OVX P4 group express markers of goblet cells (*Olfm4, Krt8, Spdef, Muc5b*). The overall lower expression of these genes in Sec cluster 1 compared to Sec cluster 2 may indicate Sec-1 cells are in an earlier stage of goblet cell differentiation. Interestingly, Sec cluster 3 which declined in the OVX P4 group expressed *Muc1* and *Krt8* but little to no expression of goblet cell markers (*Olfm4, Spdef, Muc5*b). These studies support a role of P4 in goblet cell differentiation and are consistent with prior studies that identify P4 as a key regulator of secretory lineage commitment.

Differentially expressed genes in each epithelial subtype cluster in the OVX P4 datasets as compared to OVX C were assessed (**Figure 6E**). Upregulated genes in Sec 1 and Sec 2 clusters included goblet cell markers (*Spdef, Muc5b, Clca1*) while downregulated genes were markers of squamous keratinizing cells (*Krt16, Krt6, Cnfn*). While the number of Sec 3 cells declined in the OVX P4 group, relatively few genes were differentially expressed in this cluster compared to the OVX C group. Interestingly, in basal and luminal clusters, P4 significantly upregulated genes associated with secretory cells (*Cldn10, Muc5b, Tff1, Clca1, Agr2*), while downregulating genes associated with squamous-derived luminal cells (*Krt16, Krt6b, Spink12, Mt4, Sprr2d, Dsg1a, Ifitm1, Cnfn*) (**Figure 6E**). Gene Ontology (GO) pathway analysis (**Figure 6F**) revealed progesterone-induced upregulation of glycoprotein metabolic processes and downregulation of keratinization pathways across subtypes. This is further supported by *in situ* RNA hybridization for keratinocyte genes, *Dsg1a and Krt16* **(Figure S6A).** Both genes are expressed at low levels in the OVX C group, induced in OVX E2 and suppressed in the OVX P4 treated groups. Basal cells had reduced cell migration while upregulated pathways in luminal cells and secretory cells were consistent with increased secretory cell function (**Figure 6F**). Progesterone-induced transcription factor (TF) activity analysis using SCENIC identified numerous TFs demonstrated to regulate goblet or secretory cell development or function, specifically enriched in Sec 1 and Sec 2 clusters (**Figure 6G**). Common to all clusters and consistent with known progesterone receptor (PR) functions were TFs necessary for suppression of pro-inflammatory responses **(Figure S6C).** In addition, TRIM28, previously described as a binding partner for PR necessary for uterine function, was among the top 12 TFs with high activity in the luminal cluster (43). Collectively, these findings support the role of progesterone in reorganizing cervical epithelial cell states. Specifically, progesterone directs Olfm4+ progenitors towards goblet cells differentiation, activating a transcriptional program that promotes secretory function while simultaneously downregulating the expansion and/or differentiation of Trp63+ progenitors into squamous luminal subtypes.

## DISCUSSION

The findings of this study demonstrate the adult mouse lower FRT epithelium is maintained by two progenitor pools: a columnar Olfm4+ progenitor that gives rise to secretory goblet cells, and a Tp63+ basal progenitor that gives rise to squamous keratinized luminal cells. Differentiation of these specialized luminal lineages is regulated, in part, by the steroid hormone environment. Specifically, progesterone promotes transcriptional programs that drive goblet cell differentiation from Olfm4+ progenitors, while limiting proliferation and differentiation of Trp63+ basal progenitors

The FRT is lined by secretory epithelia with regionally distinct functions. During pregnancy, secretions from the uterine epithelia support implantation and fetus development while cervical epithelia secretions form a mucosal barrier that protects against ascending infections to ensure a term pregnancy (9, 44–46). This study highlights lineage distinctions in the lower FRT compared to the uterus that are critical for the formation of a mucosal barrier. In particular, the transcriptional specification of goblet cell progenitors is established by PND 15 in the cervix and vagina. In the gastrointestinal and respiratory tracts, goblet cells arise from simple columnar progenitors (47). Similarly, in the human endocervix, goblet cells arise from columnar epithelium and are present in the endocervix before and during pregnancy (**Figure 5D**) (25, 48). Here we extend this paradigm to the mouse cervix, showing that goblet cells have a columnar origin and are derived from a luminal progenitor pool that is distinct from the basal progenitor population present in the cervix and vagina.

Prior studies support the existence of a single progenitor that maintains the adult mouse vaginal and cervical epithelium. Lineage tracing studies in adult ovariectomized mice identified a basal Axin2+ stem cell capable of regenerating all layers of the vaginal keratinized epithelium in response to E2 exposure (41). Similarly, clonal expansion studies of Trp63+ basal progenitor cells from mouse or human cervix generated both keratinized epithelia (KRT10+ or KRT13+) and secretory cells (MUC1+) (18). From these studies, the authors concluded that basal progenitors could give rise to both squamous and secretory cells. However, these prior studies did not demonstrate that goblet cells originate from basal progenitors, nor was it appreciated that goblet cells are present in the mouse vagina and ectocervix.

Collectively our findings revise the current model in which columnar epithelium lining of the mouse lower reproductive tract at birth undergoes postnatal transdifferentiation to squamous epithelium induced by the expression of the transcription factor, Trp63, on PND 5-8 (12, 13, 49, 50). We propose a refinement to this model during postnatal development (Graphical Abstract), in which a portion of Krt8+ cells acquire Trp63 expression and transition to squamous epithelium, while a subset remains Krt8+Trp63- and subsequently acquire Olfm4 expression. This Olfm4 population serves as the unipotent progenitor pool for adult goblet cells in the cervix and vagina. Thus, the cervical and vaginal epithelium in mice harbors both basal squamous Trp63+ and luminal columnar Olfm4+ progenitors.

While Olfm4 is not required for goblet cell differentiation or barrier function in mice or humans, Olfm4 serving as a progenitor marker in the murine cervix parallel findings in other epithelial tissues such as the urethral tube and prostate epithelium, where Olfm4 also has been shown to mark progenitor-like cells (28). OLFM4 expression occurs in a spatial gradient with Olfm4+ progenitors first appearing in vaginal epithelia and later extending to the ectocervix and endocervix between PND 15–25. The possibility that Olfm4 is a conserved marker for progenitor secretory cells during FRT development in the human is supported by the expression of Olfm4 in lower vaginal epithelium and urethral epithelia in single cell transcriptomic datasets from females at postconception weeks 11-21 (51). In addition, the transcriptional regulator of goblet cells, SPDEF is also expressed in this dataset in the lower vaginal epithelia, uterus/cervix epithelia, and fallopian epithelia.

Though we observed distinct populations of epithelial layers expressing Trp63 or Krt8 in the postnatal period, we cannot rule out the possibility that a small pool of bipotent progenitors persist to adulthood in the lower FRT and give rise to both the basal Trp63+ and luminal Olfm4+ progenitor pools in specific physiological periods (e.g., pregnancy) or with tissue injury or disease. This possibility is supported by the presence of minor populations of Trp63 and Krt8 co-positive cells in cervices of mice and humans. In our studies, cells co-expressing squamous (TRP63) and columnar marker (KRT8), while not uniform across all patient, resemble “reserve cells” previously described in the human cervix (52, 53) and may reflect a transitional, plastic cell state.

Overall, this study advances our understanding of lower tract epithelial differentiation during the postnatal developmental period in mice and reveals previously unrecognized species similarities between mice and humans in which goblet cell and squamous keratinized cell subtypes are derived from columnar and basal progenitors respectively. In humans, columnar progenitors are restricted to the endocervix, while basal squamous progenitors are confined to the ectocervix, potentially explaining the spatial restriction of goblet cells in humans compared with their broader distribution in mice.

This framework provides insight into epithelial plasticity required to maintain barrier homeostasis in the cycling and pregnant cervix. The rapid pivot in luminal epithelial subtypes necessary to maintain tissue homeostasis raises questions as to the cell intrinsic and extrinsic signals that regulate this process. Steroid hormones and growth factors, are key regulators of reproductive tract epithelial proliferation, differentiation and maintenance of cell-fate (19, 20, 54–56). Prior studies demonstrate region-specific signalling cues from the underlying stroma drive epithelial differentiation during postnatal development and in the adult to maintain cell-fate specification (40, 57). In addition, exogenous exposure of hormones, for example diethylstlibestlrol, during FRT development in mice or humans perturbs epithelial differentiation in the adult FRT and contributes to risk of adult-onset cancers of the FRT (15, 58) Our findings suggest that hormone signalling fine-tunes the balance between basal and columnar cervical progenitor populations. Future studies will aim to define the intrinsic and extrinsic signals governing this balance and to determine how goblet cell function contributes to mucosal defence and pregnancy outcomes.

## LIMITATIONS OF THE STUDY

Our study has several limitations. We refer to the basal progenitor as a Trp63+ basal progenitor, based on prior Trp63 gene deletion studies (12, 14). Future lineage tracing studies using isoform- specific Trp63 reporter Cres are necessary to solidify this model. Finally, human tissue analyses were limited to a small number of NP and late-pregnant samples, restricting the ability to capture dynamic changes across the menstrual cycle and gestation.

## METHODS

### Reagents and antibodies

The following chemicals were used: Tamoxifen (Sigma-Aldrich #T5648; CAS#10540-29-1), Progesterone pellets (Innovative Research of America #P-131), Estrogen pellets (Innovative Research of America #E-121). The following primary antibodies were used: Rabbit anti-P63 (D9L7L)(Cell signaling #39692S; RRID:AB_2799159), Mouse Purified anti-Cytokeratin 8 (Biolegend #904804; RRID:AB_2616821), Rabbit anti-Olfm4 mAb (D6Y5A) (Cell signaling #39141; RRID:AB_2650511), Goat tdTomato polyclonal antibody(My BioSource #MBS448092; RRID:AB_2827808), Rat anti-Ki67 mAb (SolA15)(ThermoFisher scientific #14-5698-82; RRID:AB_10854564, Mouse anti-Keratin 8( Abcam #Ab9023; RRID:AB_306948), Ghost Viability Dye 510(Tonbo Biosciences #SKU-13-0870-T100), Rat anti-mouse CD16/CD32 (BD Biosciences #553141; RRID:AB_394656), Anti-EpCAM(Abcam #ab71916; RRID:AB_1603782), Goat anti-rabbit IgG Alexa Fluor 488(ThermoFisher scientific #A11008; RRID:AB_143165), Donkey anti-goat IgG Alexa Fluor 647( #A21447; RRID:AB_141844), Goat anti-rabbit IgG Alexa Fluor 594(ThermoFisher scientific #A-11012; RRID:AB_2534079), Goat anti-Mouse IgG Alexa Fluor 488(ThermoFisher scientific #A-11001; RRID:AB_2534069), Goat anti-Rat IgG Alexa Fluor 488(ThermoFisher scientific #A-11006; RRID:AB_2534074), Donkey anti-Goat IgG Alexa Fluor 594(ThermoFisher scientific #A-11058; RRID:AB_2534105). The following Bacterial strains Escherichia (E.) coli O55:K59(B5):H-(ATCC #12014; strain CDC 5624-50 (NCTC 9701]), Escherichia (E.) coli K12 nonpathogenic strain (Gift from Dr. Michael House (Tufts University)pAKlux2.1 plasmid). The following RNAscope probe was used Mouse Spdef probe(Advanced cell diagnostics #544421), Mouse Muc5b probe(Advanced cell diagnostics #471991), Mouse Muc1 probe(Advanced cell diagnostics #421871), Mouse Olfm4 probe(Advanced cell diagnostics #311831), Mouse Dsg1a(Advanced cell diagnostics #842861), Mouse Krt16(Advanced cell diagnostics #1153201), Human spdef(Advanced cell diagnostics #543151), Human Muc5b(Advanced cell diagnostics #449881).

## EXPERIMENTAL MODEL AND SUBJECT DETAILS

### Animals

All animal experiments and protocols were approved by the Institutional Animal Care and Use Committee at UT Southwestern. All mice were housed in a controlled environment with regulated temperature and a 12-hour light/dark cycle, having free access to water and food in the animal facility. In this study, we used the wild-type C57BL6/129 (B6129F1) mouse strain, along with Olfm4-IRES-eGFPCre^ERT2^, Krt8-Cre/ERT2(17Blpn/J) (strain 017947), and Rosa26^tdTomato^ (strain 007914) mice. The Olfm4-IRES-eGFPCre^ERT2^ mice were generated in the lab of Dr. Hans Clevers and kindly provided by Dr. Linda Samuelson at the University of Michigan with permission from Dr. Clevers. The Rosa26^tdTomato^ mice were provided by Dr. Chun-Li Zhang at UTSW. The Krt8-Cre/^ERT2^(17Blpn/J) mice were sourced from Jackson Laboratories. The Olfm4-IRES-eGFPCre^ERT2^ and Krt8-Cre/^ERT2^(17Blpn/J) lines have been previously described (22, 59). OLFM4^-/-^ mice and OLFM4 WT mice were generated previously (60) and provided by Drs Wenli Liu and Griffin Rogers at NHLBI.

### Microbe strain

Two *Escherichia coli* strains were used in this study. Live *E. coli* (serotype O55; ATCC) (20 μl bacteria equal to 10^5^ CFU) was inoculated into vagina on D15, and mice were observed for 72 hours for signs of PTB as per prior study protocol (61). The bacteria were cultured aerobically at 37°C in sterile, nonselective Luria-Bertani (LB) broth overnight with shaking at 200 rpm. Cultures were harvested by centrifugation (4000g for 5 minutes) and re-suspended at the desired concentration with PBS. The correct cell concentration for vaginal inoculation was calculated by measuring an OD600 against the standard growth curve for that bacterial strain. To confirm the dose of bacteria given, an aliquot of the bacterial culture was serially diluted 1:100 to an approximate bacterial concentration of 500 CFU/ml. A 100-μl aliquot was plated under sterile conditions, and colonies were counted after overnight incubation at 37°C to confirm the accuracy of the CFU. Delivery within 72 hours of inoculation is considered preterm birth, whereas delivery on or after GD19 is considered a term birth.

Nonpathogenic *Escherichia coli* K12 used for bioluminescent imaging was constructed according to a previous protocol and sent as a gift from our collaborator Dr. Michael House (62). Bacterial inoculate was prepared by growing *E.coli* K12-lux in 30 mL LB broth containing 50 mg/mL ampicillin overnight at 37C with shaking at 200 rpm. Cultures were harvested by centrifugation. The correct cell concentration for vaginal inoculation was calculated by measuring an OD600 against the standard growth curve for that bacterial strain. After centrifugation, the bacterial pellet was resuspended in a predetermined volume of PBS and introduced to the mice. Ampicillin resistant *Escherichia coli* colonies were checked for light emission at peak emission of 490 nm (IVIS Spectrum Imaging System). Vaginal inoculation of 20 μl of 1 x 10^9^ CFU *E. Coli* was performed at GD15 followed by whole-body bioluminescence imaging at 0 and 12 hours post inoculation. At 12 hours, the mice were sacrificed and dissected. Fluorescence was detected with the IVIS Spectrum Imaging System in the amniotic sac, placenta, uterus, and pups.

### Method Details

#### Tissue collection

All mice used in NP and pregnant studies were 2-6 months old and nulliparous. The stage of the non-pregnant estrus cycle in NP females was determined by vaginal cytology, based on the presence, absence, and proportions of cornified epithelial cells, nucleated epithelial cells, and leukocytes (63). Adult mice used in this study were all females. Timed pregnancies were generated by pairing females with males overnight (4 pm to 8 am), and the following morning females were checked for the presence of a vaginal plug. The day that a plug was detected was designated as gestational day (GD) 0, with birth typically occurring on GD19. Non-pregnant (estrus, metestrus, diestrus), pregnant (GD6, 12, 15, 18, 19), and in-labor (IL) samples were collected for analysis. The Olfm4-Cre^ERT2^;Rosa26^tdTomato^ line, delivered on GD19-20. In-labor (IL) samples were collected on GD19 or 20 after visual confirmation of the delivery of 1-2 pups. For tissue collection in the postnatal period in wild type mice, cervical tissues were harvested at postnatal days (PND): PND1, PND5, and PND15, PND25 following sex determination of the pups, as previously described (64).

#### Lineage tracing studies

The Olfm4 IRES-eGFPCre^ERT2^ and Krt8 Cre/^ERT2^(17Blpn/J) mice were crossed with Rosa26^tdTomato^ mice to generate Olfm4-IRES-eGFPCre^ERT2^; Rosa26^tdTomato^ and Krt8Cre/^ERT2^; Rosa26^tdTomato^ reporter lines. Genotyping was performed via PCR on genomic DNA extracted from tail samples (65). For lineage-tracing, reporter females were paired overnight with Rosa26^tdTomato^ males (4 pm to 8 am), and vaginal plugs were checked the following morning. The day of plug detection designated GD0, with labor occurring between GD19 and 20 days. IL samples were collected on GD19/20 following visual confirmation of the delivery of 1-2 pups. To induce Cre-mediated recombination, adult females heterozygous for Olfm4-IRES-eGFPCre^ERT2^;Rosa26^tdTomato^ or Krt8 Cre/^ERT2^;Rosa26^tdTomato^ were injected intraperitoneally with a single dose of tamoxifen (Sigma, T5648) dissolved in corn oil (1 mg/g body weight in 100 µL) at 8 weeks of age. For experiments conducted in ovariectomized Olfm4-IRES-eGFPCre^ERT2^; Rosa26^tdTomato^ females, mice received two consecutive daily injections of tamoxifen (1 mg/day), then were rested for one week prior to ovariectomy as described below in the OVX Methods section.

#### Lineage tracing performed at both short-and long-term intervals

For short-term induction, tamoxifen was administered during estrus or diestrus, or on GD11, and tissues were collected at estrus, diestrus, or GD12, respectively. For long-term induction, tamoxifen was administered during non-pregnant stages, followed by timed mating. Tissues were collected during the subsequent reproductive cycle at estrus (postpartum), diestrus, and GD15 of a second pregnancy.

#### Ovariectomized (OVX) studies

C57BL6/129 female mice (7 to 8 weeks old) were OVX and allowed a 2-weeks recovery period before treatment. Females were then separated and assigned to one of the three groups: Control, estrogen (E2) and progesterone (P4). Control group: OVX-control mice received no treatment during the 15-day experimental period. Estrogen (E2) group: OVX mice were implanted with a 0.05 mg, 21-day release pellet, (Innovative Research of America) providing an estimate daily release of 2µg. Progesterone (P4) group: OVX mice received a subcutaneous injection of E2 (100 µg/d in 100 µL of corn oil) for 2 consecutive days to induce progesterone receptor expression, followed by implantation of a 25 mg, 21-day release pellet, (Innovative Research of America) providing an estimate daily release of 1mg. Tissues from all three groups were collected after 15 days of treatment for immunostaining. For single-cell RNA-seq library preparation, cervical cells were isolated from OVX control and OVX P4 groups as described previously (3, 66).

#### Mouse tissue collection and histological evaluation

Cervices and uteri were isolated by dissection at the uterocervical junction, and all vaginal tissue was removed. Cervices and uterine horns were collected for longitudinal sections with the cervical canal visible throughout each section. All tissues were fixed in 4% paraformaldehyde (Sigma-Aldrich) in 1x PBS for 24 hours at 4°C and then transferred to 1x PBS. Tissue comprising the cervix, vagina and part of the uterine horns were embedded in paraffin and 5 mm thick longitudinal sections were cut. Hematoxylin (HE) and Alcian blue-PAS staining were performed by the Histo-Pathology Core at UT Southwestern Medical Center.

#### Immunohistochemistry and Immunofluorescence staining

Immunohistochemistry (IHC) and immunofluorescence (IF) were performed according to standard protocols. IHC/IF was performed on deparaffinized and rehydrated 5 µm sections following antigen retrieval in sodium citrate buffer (10 mM, pH 6) for 20 min at 4°C, as previously described (3). Staining was repeated on at least three sections from three independent animals (biological replicates). Representative immunostainings are shown in the manuscript.

#### In situ mRNA hybridization

RNAscope was performed using RNAscope® Multiplex Fluorescent Reagent Kit V2 (Advanced Cell Diagnostics) as previously described (3). RNAScope was repeated on tissue sections collected from two to three biological replicates. Representative images were included in the manuscript.

#### Human Tissues

All experimental procedures involving human tissues were approved by the Institutional Review Board at the University of Texas Southwestern Medical Center (IRB# 112010-047). Nonpregnant (premenopausal) and pregnant (34–36 weeks gestation) cervical tissue were obtained from the UTSW Department of Obstetrics and Gynecology Tissue Biorepository. Pregnant ectocervical and endocervical samples were collected at the time of hysterectomy for complications associated with placenta accreta; cases with placental invasion near the cervix were excluded. Transmural sections (10 × 10 mm) were wrapped in Kimwipes to preserve endocervical canal epithelial cells and fixed in 4% paraformaldehyde for 24 hrs, followed by transfer to PBS. Longitudinal sections (5 mm thick) of endocervix or ectocervix were then prepared. Hematoxylin and eosin (H&E) staining was performed by the Histo-Pathology Core at UTSW.

#### Systemic LPS-Induced PTB

Preterm labor was induced by intraperitoneal injection of 100 µL lipopolysaccharide (LPS; serotype O55, Sigma-Aldrich) at 5mg/kg concentration in GD15 in wild-type and OLFM4-deficient mice. Animals were either observed for preterm birth or sacrificed 6 hrs after injection (prior to onset of preterm birth) for tissue collection.

#### Image analysis

The whole-slides immunohistochemistry samples were scanned using the Hamamatsu Nanozoomer S60 and viewed using NDP View2 software, as previously described (**3**). Immunofluorescence and RNAscope images of longitudinal cervical sections were acquired using Zeiss Axioscan Z1 digital slide scanner with ZEN 3.0 (software) at 20x magnifications. To ensure consistency across time points, all fluorescent images, including RNAscope with TSA dyes, were captured under identical detector gain and laser power settings using the appropriate filter configurations. Standard excitation and emission settings were DAPI (358/461 nm), FITC/TSA 520 (494/525 nm), Cy3/TSA 570 (550/570 nm) and Cy5/TSA 690 (676/694 nm). For temporal comparisons, optimal imaging conditions determined at the lowest-signal time point were applied to all subsequent time points, and images were linearly rescaled per channel. Scale bars were added using Image J.

#### Cell isolation for tdTomato sorting from Ai14Olfm4 lineage mice

Olfm4-IRES-eGFPCre^ERT2^;Rosa26^tdTomato^ females (n = 3; two independent experiments) were administered two consecutive intraperitoneal doses of tamoxifen (1 mg each) during the non-pregnant stage. Cervical tissues were harvested at diestrus, minced (∼1 mm³), and digested overnight at 4°C on a rotating platform in Dispase 5 U/ml (Stem cell tech, 7913) solution. Digests were diluted in ice-cold PBS or Advanced DMEM/F12 supplemented with GlutaMax, HEPES, and Penicillin/Streptomycin, filtered through a 40 µm strainer, and centrifuged (1500 rpm, 5 min, 4°C). Pellets were resuspended in cold PBS, and supernatants were reduced to ∼100–200 µL for flow cytometry.

#### Cell Staining for Flow Cytometry and Analysis

Single-cell suspensions were divided into experimental and control tubes. Controls included: (i) unstained wild-type NP cells (autofluorescence baseline), (ii) viability dye only, (iii) single-color controls for EpCAM and tdTomato, and (iv) secondary antibody only controls (488 and 647 nm). Experimental samples were incubated with Ghost Viability Dye 510 (Tonbo Biosciences; Cytek SKU-13-0870-T100), followed by Fc receptor blockade with BD Pharmingen™ anti-CD16/CD32 (Mouse BD Fc Block™, Cat#553141). Because tdTomato reporter expression in the NP diestrus cervix was low and not reliably detected by flow cytometry, antibody-based detection was used to enhance signal. Cells were incubated on ice for 30 min with rabbit anti-EpCAM (Abcam, ab71916) and goat anti-tdTomato (My Biosource, MBS448092). After washing, cells were incubated on ice for 30 min with Alexa Fluor 488 conjugated goat anti-rabbit IgG (Invitrogen, A11008) and Alexa Fluor 647conjugated donkey anti-goat IgG (Invitrogen, A21447). Final washes were performed, and cells were resuspended in Advanced DMEM/F12 supplemented with GlutaMax, HEPES, and Penicillin/Streptomycin and passed through 30 µm-filtered FACS tubes. Flow cytometry and sorting were performed using a Cytek Aurora spectral cytometer. Live epithelial cells were gated by EpCAM expression and viability dye exclusion. In Ai14-Olfm4 samples, luminal progenitors were defined as EpCAM⁺ tdTomato⁺, while basal progenitors were defined as EpCAM⁺ tdTomato⁻. Data were analyzed using FlowJo_v10.9.0.

#### Organoid Culture

Flow-sorted populations (EpCAM⁺ tdTomato⁺ luminal progenitors and EpCAM⁺ tdTomato⁻ basal progenitors) were embedded separately in growth factor reduced basement membrane extract (BME, Cultrex) and cultured in cervical organoid expansion medium as previously described (**67**). Organoids were maintained at 37 °C in 5% CO₂, with medium replaced every 3 days. After 10–12 days, organoids were imaged by confocal microscopy to verify tdTomato⁺ luminal cell contribution. Organoids from 3 biological replicates were maintained for at least three passages over approximately 5 weeks (passages every 12 days). Organoid formation assays were conducted for each biological replicate using three technical (wells) replicates (5000 cells/25 μL drops; 2 drops per well) and the data are presented as the mean ± SEM. Statistical differences between two groups were determined with Welch’s tLJtest (GraphPad Prism 10.5), and statistical significance was defined as *p* < 0.05.

#### RNA isolation and quantitative real time PCR

Total RNA was extracted from tissues using RNA-STAT 60 (Tel-Test) and treated with DNase I (DNA-Free, Ambion). cDNA was synthesized from 1 µg of total RNA in a 20 µL reaction using the iScript Reverse Transcription Supermix (Bio-Rad). Quantitative real-time PCR was performed with SYBR Green chemistry on a PRISM 7900HT Sequence Detection System (Applied Biosystems). Reactions were run in triplicate with 20 ng cDNA per well. Gene expression levels were normalized to the housekeeping gene 36B4, and relative expression was calculated using the 2–ΔΔCt method. Cell junction gene expression in cervical tissues was assessed using the RT² Profiler PCR Array Mouse Cell Junction Pathway Finder Kit (Qiagen) according to the manufacturer’s instructions. Predesigned SYBR Green primers for mouse inflammatory genes (TNFα, IL6, IL1β, and CXCL2) were purchased from Sigma-Aldrich (St. Louis, MO). The following probes were used for qPCR IL6 Forward (5’-TCG TGG AAA TGA GAA AAG AGT TG -3’), IL6 Reverse (5’- AGT GCA TCA TCG TTG TTC ATA CA -3’), Cxcl2 Forward (5’-GAA CAT CCA GAG CTT GAG TGT GA -3’), Cxcl2 Reverse (5’-CCT TGA GAG TGG CTA TGA CTT CTG T -3’), IL-1β Forward (5’-GCC CAT CCT CTG TGA CTC ATG -3’), IL-1β Reverse (5’- AGC CTG TAG TGC AGC TGT CTA ATG -3’), TNF-α Forward (5’- CTG AGG TCA ATC TGC CCA AGT AC -3’), TNF-α Reverse (5’- CTT CAC AGA GCA ATG ACT CCA AAG -3’).

#### Single cell RNA-sequencing of OVX samples and analysis

Reads were aligned to the mouse genome (mm10) using Cell Ranger v3.0.2. Downstream processing was performed with Seurat v5 (68) following the standard workflow (https://satijalab.org/seurat/articles/get_started_v5_new). Cells with >300 detected genes and <10% mitochondrial transcripts were retained; genes detected in at least 10 cells were included. After filtering, 4,148 and 7,659 cells were retained for the OVX control and OVX P4 samples, respectively. Normalization across conditions was performed using SCTransform, and datasets were integrated with the Harmony algorithm via the Seurat “integrate layers” function without regression of variables. The top 30 principal components were used for clustering with a resolution of 0.8, yielding 21 clusters. Cell types were assigned based on canonical marker genes (3), resulting in five major cell types. The epithelial cluster was subsetted for further analysis of epithelial subtypes using established markers (3). Differential gene expressions were assessed between OVX Control and OVX P4 samples by down-sampling to the smallest group size and applying the Seurat *FindMarkers* function. To evaluate transcription factor activity at single-cell resolution, the SCENIC workflow (69) was applied, and differential activity was quantified using AUCell scores analyzed with the *limma* package (70).

#### Quantification and statistical analysis

Statistical analyses were performed using standard parametric and non-parametric tests as appropriate for each dataset. For murine inflammatory gene expression studies, relative mRNA levels of IL6, CXCL2, IL1β, and TNFα in uterine and cervical tissues from WT and Spdef knockout mice exposed to intraperitoneal lipopolysaccharide (LPS; 5 mg/kg, 6 hours) versus PBS control (n = 5 per condition) were compared using unpaired t-tests, with a significance threshold of p < 0.05. Preterm birth outcomes following vaginal inoculation with live pathogenic *E. coli* (10LJ CFU) , intraperitoneal LPS administration (5 mg/kg) were compared between WT and knockout mice using Fisher’s exact test. *E. coli*–induced preterm birth (p = 0.20) in WT and knockout mice were compared using the unpaired t-test (p = 0.94).

## Data availability

OVX and OVX P4 single cell datasets have been deposited at GEO and are publicly available as of the date of publication. The accession number for the scRNA-seq data reported in this paper is GEO: GSE322687.

## ACKNOWLEDGEMENTS

This work was supported by the Burroughs Welcome Fund grant (1019804) and the National Institute of Health (R01HD110147) to M.M. Computational resources were provided by the BioHPC supercomputing facility located in the Lyda Hill Department of Bioinformatics, UT Southwestern Medical Center, URL: https://portal.biohpc.swmed.edu. We would like to acknowledge the assistance of the UT Southwestern Quantitative Light Microscopy Core, a Shared Resource of the Harold C. Simmons Cancer Center, supported in part by an NCI Cancer Center Support Grant, 1P30 CA142543-01. We also acknowledge and thank Philip Dryden in the laboratory of Dr. Tiffany Reese in the UT Southwestern Department of Immunology for assistance with IVIS bioluminescence imaging experiments. We thank the UT Southwestern Medical Center Whole Brain Microscopy Facility (RRID: SCR_017949) for assistance with whole slide imaging. We thank Lily Farid for help with immunohistochemistry. We thank Dr. Andrew Kelleher at the Univ of Missouri for reading the manuscript and for valuable discussions. The graphical abstract was created by Angela Diehl.

## AUTHOR CONTRIBUTIONS

S.P.M, Y.F., and M.M. designed the experiments. S.P.M. performed experiments and data analysis for all lineage tracing studies, OVX studies and spatial analysis of human cervix. Y.F and E.P. conducted experiments and data analysis in the Olfm4 knockout mice. G.B. performed computational analysis of OVX datasets. L.W. generated scRNAseq libraries. J.W. assisted in generating timed pregnant mice for the described studies and tissue collection. M.F.R. provided human cervical tissues for described studies. S.P.M, Y.F., and M.M. wrote the first draft of the paper. All authors read and provided edits to the manuscript. M.M. and G.C.H. secured funding to support this project and provided intellectual support for all aspects of the work.

## REFERENCES

1. Ueda Y, Mogami H, Chigusa Y, Kawamura Y, Inohaya A, Takakura M, et al. Hyposecretion of cervical MUC5B is related to preterm birth in pregnant women after cervical excisional surgery. Am J Reprod Immunol. 2024;91(3):e13832.

2. Lacroix G, Gouyer V, Rocher M, Gottrand F, Desseyn JL. A porous cervical mucus plug leads to preterm birth induced by experimental vaginal infection in mice. iScience. 2022;25(7):104526.

3. Cooley A, Madhukaran S, Stroebele E, Colon Caraballo M, Wang L, Akgul Y, et al. Dynamic states of cervical epithelia during pregnancy and epithelial barrier disruption. iScience. 2023;26(2):105953.

4. Asada R, Saito A, Kawasaki N, Kanemoto S, Iwamoto H, Oki M, et al. The endoplasmic reticulum stress transducer OASIS is involved in the terminal differentiation of goblet cells in the large intestine. J Biol Chem. 2012;287(11):8144–53.

5. Chen G, Korfhagen TR, Xu Y, Kitzmiller J, Wert SE, Maeda Y, et al. SPDEF is required for mouse pulmonary goblet cell differentiation and regulates a network of genes associated with mucus production. J Clin Invest. 2009;119(10):2914–24.

6. Noah TK, Kazanjian A, Whitsett J, Shroyer NF. SAM pointed domain ETS factor (SPDEF) regulates terminal differentiation and maturation of intestinal goblet cells. Exp Cell Res. 2010;316(3):452–65.

7. Kim YS, Ho SB. Intestinal goblet cells and mucins in health and disease: recent insights and progress. Curr Gastroenterol Rep. 2010;12(5):319–30.

8. Nystrom EEL, Martinez-Abad B, Arike L, Birchenough GMH, Nonnecke EB, Castillo PA, et al. An intercrypt subpopulation of goblet cells is essential for colonic mucus barrier function. Science. 2021;372(6539).

9. Cunha GR, Kurita T, Cao M, Shen J, Robboy S, Baskin L. Molecular mechanisms of development of the human fetal female reproductive tract. Differentiation. 2017;97:54–72.

10. Robboy SJ, Kurita T, Baskin L, Cunha GR. New insights into human female reproductive tract development. Differentiation. 2017;97:9–22.

11. Terakawa J, Serna VA, Nair DM, Sato S, Kawakami K, Radovick S, et al. SIX1 cooperates with RUNX1 and SMAD4 in cell fate commitment of Mullerian duct epithelium. Cell Death Differ. 2020;27(12):3307–20.

12. Kurita T, Cunha GR, Robboy SJ, Mills AA, Medina RT. Differential expression of p63 isoforms in female reproductive organs. Mech Dev. 2005;122(9):1043–55.

13. Kurita T. Developmental origin of vaginal epithelium. Differentiation. 2010;80(2-3):99–105.

14. Romano RA, Smalley K, Magraw C, Serna VA, Kurita T, Raghavan S, et al. DeltaNp63 knockout mice reveal its indispensable role as a master regulator of epithelial development and differentiation. Development. 2012;139(4):772–82.

15. Kurita T, Mills AA, Cunha GR. Roles of p63 in the diethylstilbestrol-induced cervicovaginal adenosis. Development. 2004;131(7):1639–49.

16. Cunha GR, Sinclair A, Ricke WA, Robboy SJ, Cao M, Baskin LS. Reproductive tract biology: Of mice and men. Differentiation. 2019;110:49–63.

17. Jin S. Bipotent stem cells support the cyclical regeneration of endometrial epithelium of the murine uterus. Proc Natl Acad Sci U S A. 2019;116(14):6848–57.

18. Zhao Z, Wang Y, Wu Y, Li D, Zhang T, Ma Y, et al. Single-cell analysis defines the lineage plasticity of stem cells in cervix epithelium. Cell Regen. 2021;10(1):36.

19. Yin Y, Haller M, Goldinger L, Bharadwaj S, So E, Robles-Pinos V, et al. Retinoic acid antagonizes estrogen signaling to maintain adult uterine cell fate. Proc Natl Acad Sci U S A. 2025;122(5):e2416089122.

20. Rizo JA, Davenport KM, Winuthayanon W, Spencer TE, Kelleher AM. Estrogen receptor alpha regulates uterine epithelial lineage specification and homeostasis. iScience. 2023;26(9):107568.

21. Parks SE, Tang S, Unser AC, Bhonsley AL, Chung EM, Jiang P, et al. TGFBR2 coordinates the endometrial response to estrogen, regulating endometrial hyperplasia and fertility. Proc Natl Acad Sci U S A. 2025;122(49):e2518507122.

22. Schuijers J, van der Flier LG, van Es J, Clevers H. Robust cre-mediated recombination in small intestinal stem cells utilizing the olfm4 locus. Stem Cell Reports. 2014;3(2):234–41.

23. Forsberg JG. Cervicovaginal epithelium: its origin and development. Am J Obstet Gynecol. 1973;115(7):1025–43.

24. Chateau D, Boehm N. Regulation of differentiation and keratin 10 expression by all-trans retinoic acid during the estrous cycle in the rat vaginal epithelium. Cell Tissue Res. 1996;284(3):373–81.

25. Ueda Y, Mogami H, Kawamura Y, Takakura M, Inohaya A, Yasuda E, et al. Cervical MUC5B and MUC5AC are Barriers to Ascending Pathogens During Pregnancy. J Clin Endocrinol Metab. 2022;107(11):3010–21.

26. van der Flier LG, Haegebarth A, Stange DE, van de Wetering M, Clevers H. OLFM4 is a robust marker for stem cells in human intestine and marks a subset of colorectal cancer cells. Gastroenterology. 2009;137(1):15–7.

27. Sakahara M, Okamoto T, Srivastava U, Natsume Y, Yamanaka H, Suzuki Y, et al. Paneth-like cells produced from OLFM4(+) stem cells support OLFM4(+) stem cell growth in advanced colorectal cancer. Commun Biol. 2024;7(1):27.

28. Li H, Chaitankar V, Zhu J, Chin K, Liu W, Pirooznia M, et al. Olfactomedin 4 mediation of prostate stem/progenitor-like cell proliferation and differentiation via MYC. Sci Rep. 2020;10(1):21924.

29. Spencer TE, Lowke MT, Davenport KM, Dhakal P, Kelleher AM. Single-cell insights into epithelial morphogenesis in the neonatal mouse uterus. Proc Natl Acad Sci U S A. 2023;120(49):e2316410120.

30. Gersemann M, Becker S, Nuding S, Antoni L, Ott G, Fritz P, et al. Olfactomedin-4 is a glycoprotein secreted into mucus in active IBD. J Crohns Colitis. 2012;6(4):425–34.

31. Liu W, Rodgers GP. Olfactomedin 4 expression and functions in innate immunity, inflammation, and cancer. Cancer Metastasis Rev. 2016;35(2):201–12.

32. Al Gharaibeh FN, Kempton KM, Alder MN. Olfactomedin-4-Positive Neutrophils in Neonates: Link to Systemic Inflammation and Bronchopulmonary Dysplasia. Neonatology. 2023;120(1):40–8.

33. Clemmensen SN, Bohr CT, Rorvig S, Glenthoj A, Mora-Jensen H, Cramer EP, et al. Olfactomedin 4 defines a subset of human neutrophils. J Leukoc Biol. 2012;91(3):495–500.

34. Alder MN, Mallela J, Opoka AM, Lahni P, Hildeman DA, Wong HR. Olfactomedin 4 marks a subset of neutrophils in mice. Innate Immun. 2019;25(1):22–33.

35. Liu W, Yan M, Sugui JA, Li H, Xu C, Joo J, et al. Olfm4 deletion enhances defense against Staphylococcus aureus in chronic granulomatous disease. J Clin Invest. 2013;123(9):3751–5.

36. Gipson IK, Spurr-Michaud S, Moccia R, Zhan Q, Toribara N, Ho SB, et al. MUC4 and MUC5B transcripts are the prevalent mucin messenger ribonucleic acids of the human endocervix. Biol Reprod. 1999;60(1):58–64.

37. Gipson IK. Mucins of the human endocervix. Front Biosci. 2001;6:D1245–55.

38. Arslan SY, Yu Y, Burdette JE, Pavone ME, Hope TJ, Woodruff TK, et al. Novel three dimensional human endocervix cultures respond to 28-day hormone treatment. Endocrinology. 2015;156(4):1602–9.

39. Greening DW, Nguyen HP, Evans J, Simpson RJ, Salamonsen LA. Modulating the endometrial epithelial proteome and secretome in preparation for pregnancy: The role of ovarian steroid and pregnancy hormones. J Proteomics. 2016;144:99–112.

40. Kurita T, Cooke PS, Cunha GR. Epithelial-stromal tissue interaction in paramesonephric (Mullerian) epithelial differentiation. Dev Biol. 2001;240(1):194–211.

41. Ali A, Syed SM, Jamaluddin MFB, Colino-Sanguino Y, Gallego-Ortega D, Tanwar PS. Cell Lineage Tracing Identifies Hormone-Regulated and Wnt-Responsive Vaginal Epithelial Stem Cells. Cell Rep. 2020;30(5):1463–77 e7.

42. Buchanan DL, Kurita T, Taylor JA, Lubahn DB, Cunha GR, Cooke PS. Role of stromal and epithelial estrogen receptors in vaginal epithelial proliferation, stratification, and cornification. Endocrinology. 1998;139(10):4345–52.

43. Li R, Wang T, Marquardt RM, Lydon JP, Wu SP, DeMayo FJ. TRIM28 modulates nuclear receptor signaling to regulate uterine function. Nat Commun. 2023;14(1):4605.

44. Wira CR, Fahey JV, Ghosh M, Patel MV, Hickey DK, Ochiel DO. Sex hormone regulation of innate immunity in the female reproductive tract: the role of epithelial cells in balancing reproductive potential with protection against sexually transmitted pathogens. Am J Reprod Immunol. 2010;63(6):544–65.

45. Lacroix G, Gouyer V, Gottrand F, Desseyn JL. The Cervicovaginal Mucus Barrier. Int J Mol Sci. 2020;21(21).

46. Kelleher AM, DeMayo FJ, Spencer TE. Uterine Glands: Developmental Biology and Functional Roles in Pregnancy. Endocr Rev. 2019;40(5):1424–45.

47. Gustafsson JK, Hansson GC. Immune Regulation of Goblet Cell and Mucus Functions in Health and Disease. Annu Rev Immunol. 2025;43(1):169–89.

48. Madhukaran S, Fomina YY, Mahendroo M. Cervical function in pregnancy and disease: new insights from single-cell analysis. Am J Obstet Gynecol. 2025;232(4S):S81–S94.

49. Kurita T, Cunha GR. Roles of p63 in differentiation of Mullerian duct epithelial cells. Ann N Y Acad Sci. 2001;948:9–12.

50. Terakawa J, Rocchi A, Serna VA, Bottinger EP, Graff JM, Kurita T. FGFR2IIIb-MAPK Activity Is Required for Epithelial Cell Fate Decision in the Lower Mullerian Duct. Mol Endocrinol. 2016;30(7):783–95.

51. Lorenzi V, Icoresi-Mazzeo C, Cassie C, Yayon N, Ruiz-Morales ER, Sancho-Serra C, et al. Spatiotemporal cellular map of the developing human reproductive tract. Nature. 2025.

52. Martens JE, Smedts F, van Muyden RC, Schoots C, Helmerhorst TJ, Hopman A, et al. Reserve cells in human uterine cervical epithelium are derived from mullerian epithelium at midgestational age. Int J Gynecol Pathol. 2007;26(4):463–8.

53. Martens JE, Smedts FM, Ploeger D, Helmerhorst TJ, Ramaekers FC, Arends JW, et al. Distribution pattern and marker profile show two subpopulations of reserve cells in the endocervical canal. Int J Gynecol Pathol. 2009;28(4):381–8.

54. Kurita T, Lee KJ, Cooke PS, Taylor JA, Lubahn DB, Cunha GR. Paracrine regulation of epithelial progesterone receptor by estradiol in the mouse female reproductive tract. Biol Reprod. 2000;62(4):821–30.

55. Mehta FF, Son J, Hewitt SC, Jang E, Lydon JP, Korach KS, et al. Distinct functions and regulation of epithelial progesterone receptor in the mouse cervix, vagina, and uterus. Oncotarget. 2016;7(14):17455–67.

56. Chumduri C, Gurumurthy RK, Berger H, Dietrich O, Kumar N, Koster S, et al. Opposing Wnt signals regulate cervical squamocolumnar homeostasis and emergence of metaplasia. Nat Cell Biol. 2021;23(2):184–97.

57. Cunha GR. Epithelial-stromal interactions in development of the urogenital tract. Int Rev Cytol. 1976;47:137–94.

58. Padilla-Banks E, Jefferson WN, Papas BN, Suen AA, Xu X, Carreon DV, et al. Developmental estrogen exposure in mice disrupts uterine epithelial cell differentiation and causes adenocarcinoma via Wnt/beta-catenin and PI3K/AKT signaling. PLoS Biol. 2023;21(10):e3002334.

59. Van Keymeulen A, Rocha AS, Ousset M, Beck B, Bouvencourt G, Rock J, et al. Distinct stem cells contribute to mammary gland development and maintenance. Nature. 2011;479(7372):189–93.

60. Liu W, Yan M, Liu Y, Wang R, Li C, Deng C, et al. Olfactomedin 4 down-regulates innate immunity against Helicobacter pylori infection. Proc Natl Acad Sci U S A. 2010;107(24):11056–61.

61. Akgul Y, Word RA, Ensign LM, Yamaguchi Y, Lydon J, Hanes J, et al. Hyaluronan in cervical epithelia protects against infection-mediated preterm birth. J Clin Invest. 2014;124(12):5481–9.

62. Zhang Y, Edwards SA, House M. Cerclage prevents ascending intrauterine infection in pregnant mice. Am J Obstet Gynecol. 2024;230(5):555 e1– e8.

63. Byers SL, Wiles MV, Dunn SL, Taft RA. Mouse estrous cycle identification tool and images. PLoS One. 2012;7(4):e35538.

64. Papaioannou VE, Behringer RR. Sex Genotyping Mice by Polymerase Chain Reaction. Cold Spring Harb Protoc. 2024;2024(1):108062.

65. Jacquot S, Chartoire N, Piguet F, Herault Y, Pavlovic G. Optimizing PCR for Mouse Genotyping: Recommendations for Reliable, Rapid, Cost Effective, Robust and Adaptable to High-Throughput Genotyping Protocol for Any Type of Mutation. Curr Protoc Mouse Biol. 2019;9(4):e65.

66. Madhukaran S, Hon GC, Mahendroo M. Protocol to dissociate epithelia from non-pregnant and pregnant mouse cervical tissue for single-cell RNA-sequencing. STAR Protoc. 2023;4(4):102631.

67. Lohmussaar K, Oka R, Espejo Valle-Inclan J, Smits MHH, Wardak H, Korving J, et al. Patient-derived organoids model cervical tissue dynamics and viral oncogenesis in cervical cancer. Cell Stem Cell. 2021;28(8):1380–96 e6.

68. Hao Y, Stuart T, Kowalski MH, Choudhary S, Hoffman P, Hartman A, et al. Dictionary learning for integrative, multimodal and scalable single-cell analysis. Nat Biotechnol. 2024;42(2):293–304.

69. Aibar S, Gonzalez-Blas CB, Moerman T, Huynh-Thu VA, Imrichova H, Hulselmans G, et al. SCENIC: single-cell regulatory network inference and clustering. Nat Methods. 2017;14(11):1083–6.

70. Ritchie ME, Phipson B, Wu D, Hu Y, Law CW, Shi W, et al. limma powers differential expression analyses for RNA-sequencing and microarray studies. Nucleic Acids Res. 2015;43(7):e47.

