## Supplemental Figures for "Olfactomedin4 marks luminal progenitor cells that give rise to secretory cell lineage in the mouse cervix": Supplement_Figures_Tables_Development.pdf

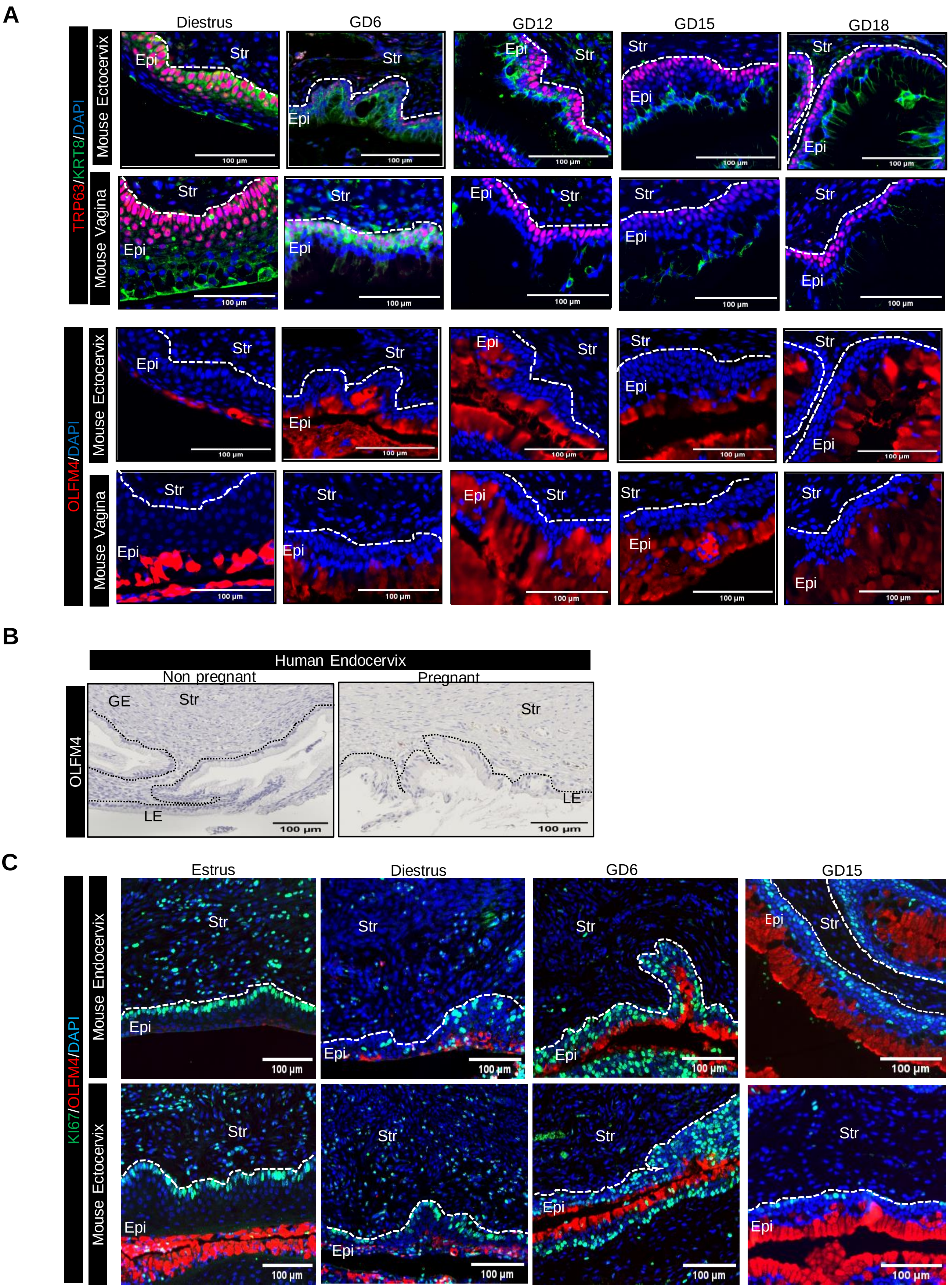

A
TRP63/KRT8/DAPI
Mouse Ectocervix
Mouse Vagina
Diestrus
GD6
GD12
GD15
GD18
Epi
Str
100 μm
OLFM4/DAPI
B
Human Endocervix
Non pregnant
Pregnant
OLFM4
GE
LE
C
KI67/OLFM4/DAPI
Mouse Endocervix
Estrus

**Supplemental Figure 1: Distinct epithelial subtypes express squamous (TRP63) or columnar (KRT8/OLFM4) markers in the ectocervix and vagina of NP and pregnant mice.**

A. Immunofluorescence staining on transverse sections of the ectocervix and vagina across non-pregnant (diestrus), pregnant (GD6, GD12, GD15, GD18) timepoints. Top panels show TRP63 (red), KRT8 (green), and nuclei (blue); bottom panels show OLFM4 (red) and nuclei (blue), illustrating temporal and spatial expression across the same timepoints. Images are representative of n = 3 biological replicates per time point, and for each marker combination the endocervix (**Figure 1**), ectocervix and vagina images in **Figure S1** are taken from the same tissue section. Scale bars = 100 µm.

B. Immunohistochemical staining for OLFM4 in human endocervix. N= 2 biological replicates per group. Scale bar 100 µm. LE: luminal epithelia, GE: glandular epithelia, Str:Stroma.

C. Immunofluorescence staining for KI67 (green), OLFM4 (red), and DAPI (blue) in the mouse cervix at estrus, diestrus, GD6, and GD15 timepoints. Images show endocervix (upper rows) and ectocervix (lower rows) with endo and ectocervical images taken from the same tissue section. n=3 biological replicates per time point and scale bars: 100 µm

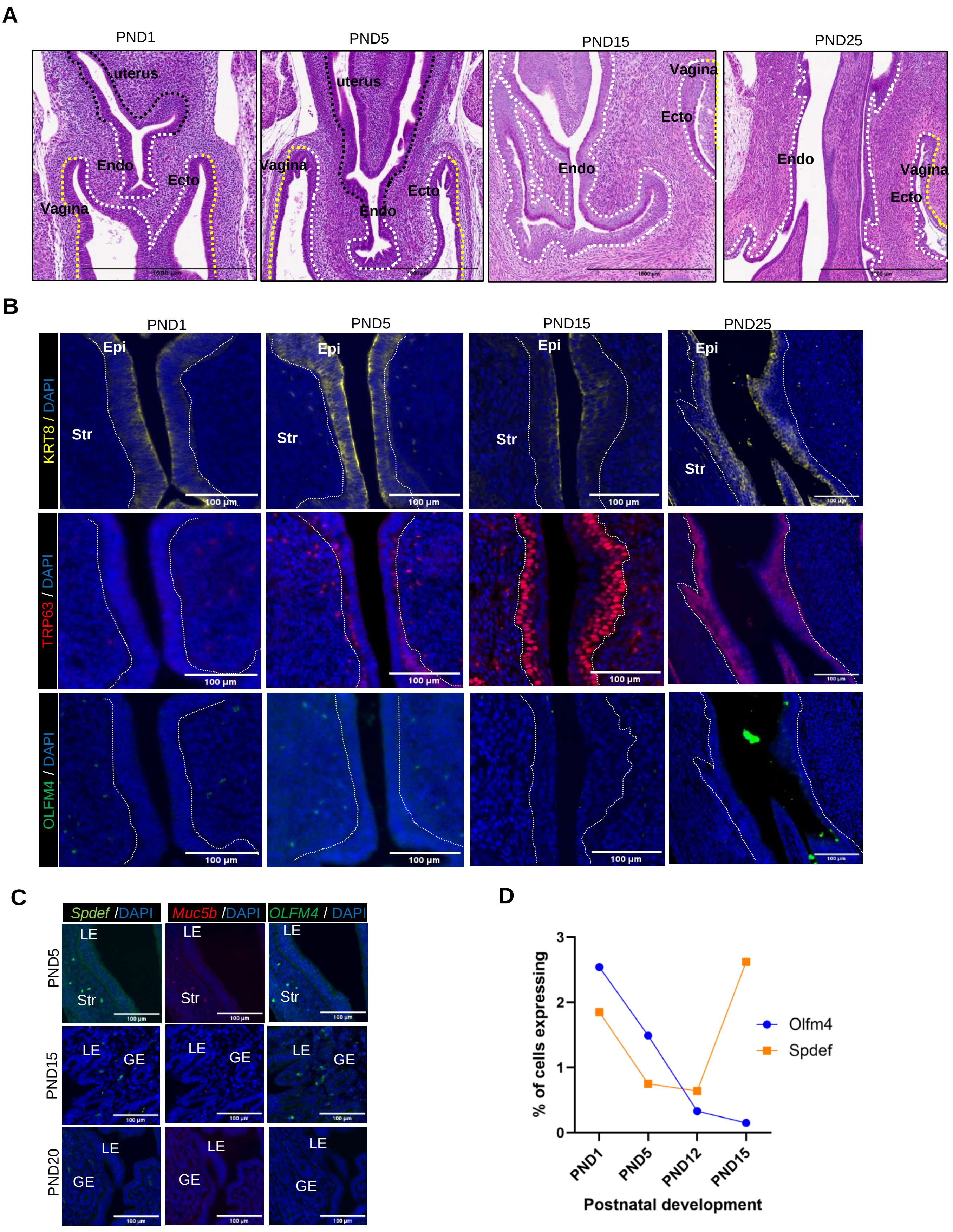
A
PND1
PND5
PND15
PND25
uterus
Endo
Ecto
Vagina
B
Epi
Str
KRT8 / DAPI
TRP63 / DAPI
OLFM4 / DAPI
100 µm
C
Spdef /DAPI
Muc5b /DAPI
OLFM4 / DAPI
PND5
PND15
PND20
LE
GE
Str
100 µm
D
% of cells expressing
Postnatal development
PND1
PND5
PND12
PND15
Olfm4
Spdef
0
1
2
3

**Supplemental Figure 2: H&E and RNA expression in the mouse female reproductive tract during development.**

A. Hematoxylin and eosin (H&E) staining of FRT at postnatal day (PND) 1, 5, 15, and 25. Anatomical compartments are outlined with colored dotted lines: cervix (white), and vagina (yellow). Scale bars = 1000 µm. Ecto, ectocervix; Endo, endocervix.

B. Magnified images taken from images in Figure 2A

C. Spdef, Muc5b and Olfm4 are not expressed in the uterus during the postnatal developmental period. Magnified images are taken from the same sections as shown in Figure 2B. LE: Luminal epithelia; GE: Glandular epithelia. scale bars: 100 µm

D. Line graph showing the percentage of cells expressing Olfm4 and Spdef in PND 1,5,12,15 uterine tissue as described in (29).

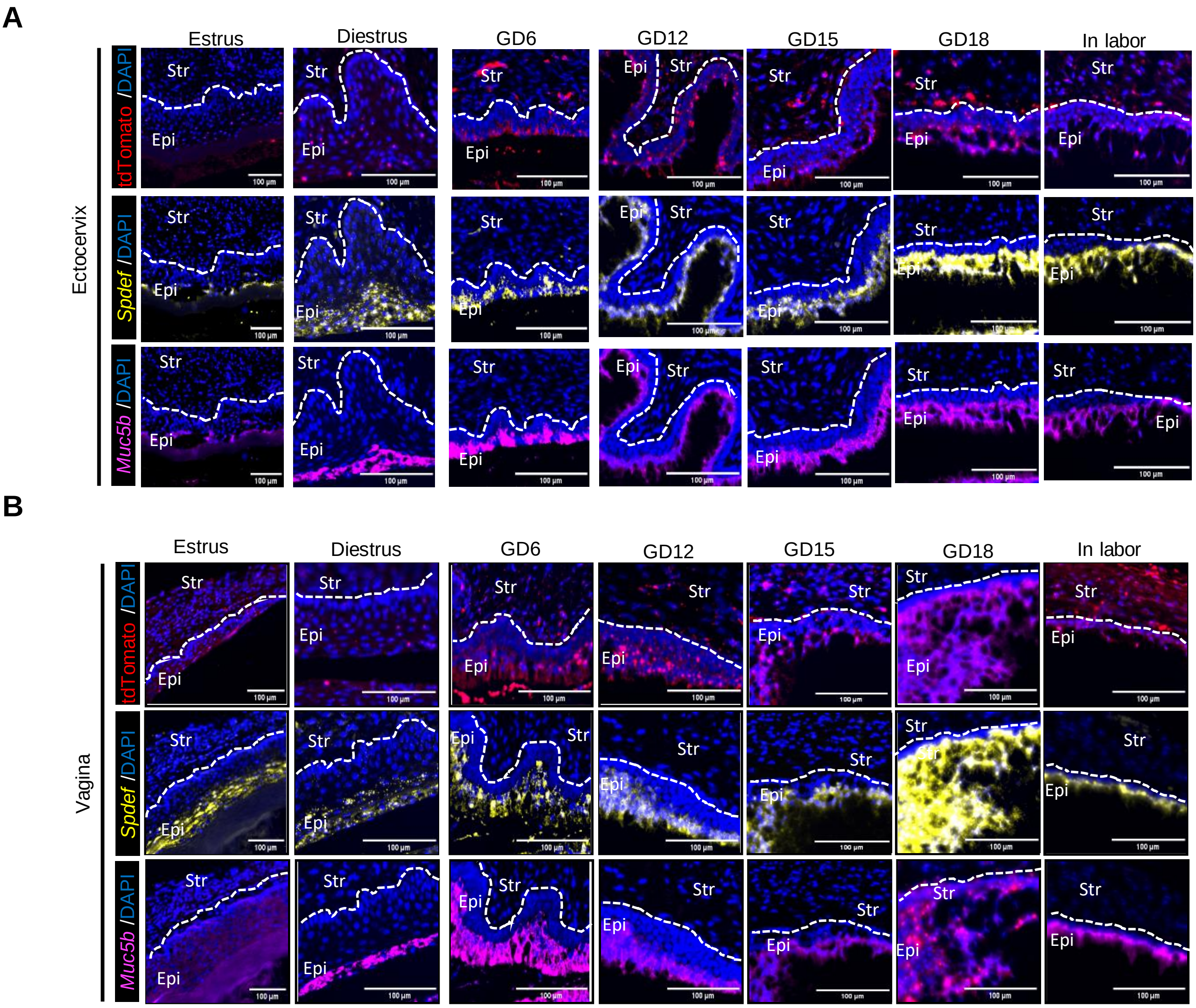

A
Estrus
Diestrus
GD6
GD12
GD15
GD18
In labor
Ectocervix
tdTomato /DAPI
Spdef /DAPI
Muc5b /DAPI
Str
Epi
100 μm
B
Vagina

**Supplemental Figure 3: Krt8$^+$ progenitors give rise to goblet cells in the ectocervix and vagina.**

A. RNAscope staining in the ectocervix for tdTomato (red, top panel), Spdef (yellow, middle panel), and Muc5b (magenta, bottom panel) across estrus, diestrus, pregnancy stages and in labor. Spdef and Muc5b were co-stained on the same tissue section and are shown in separate panels, whereas tdTomato was stained on a second section from the same animal. Images are representative of n=2 biological replicates per time points. Scale bars =100 µm.

B. RNAscope staining in the vagina for tdTomato (red, top panel), Spdef (yellow, middle panel), and Muc5b (magenta, bottom panel) across estrus, diestrus, pregnancy stages and in labor. Spdef and Muc5b were co-stained on the same tissue section and are shown in separate panels, whereas tdTomato was stained on separate section. Images are representative of n=2 biological replicates per time points. Scale bars =100 µm.

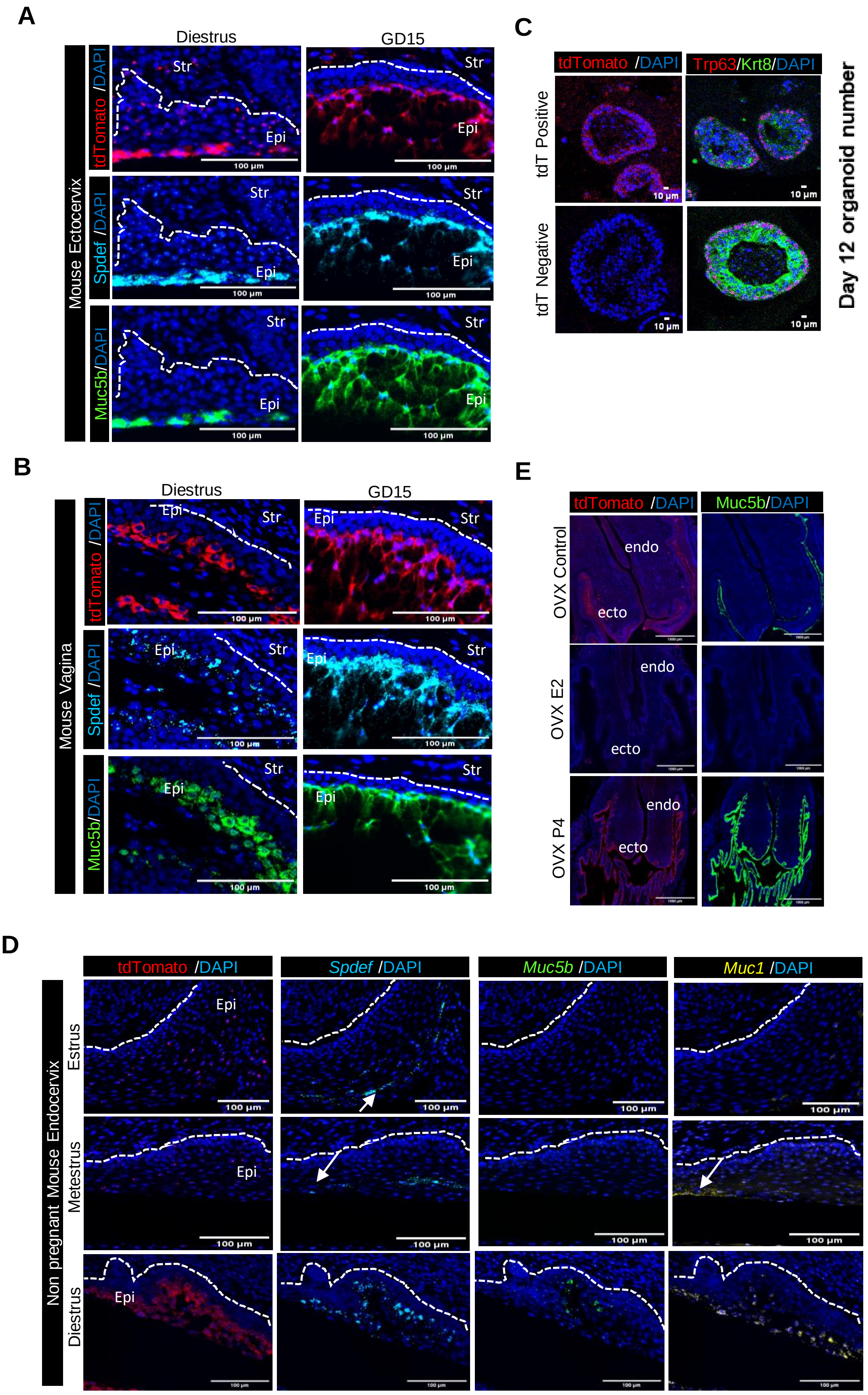

A
Diestrus
GD15
Mouse Ectocervix
tdTomato /DAPI
Spdef /DAPI
Muc5b/DAPI
Str
Epi
100 µm
B
Diestrus
GD15
Mouse Vagina
tdTomato /DAPI
Spdef /DAPI
Muc5b/DAPI
Str
Epi
100 µm
C
tdTomato /DAPI
Trp63/Krt8/DAPI
tdT Positive
tdT Negative
10 µm
Day 12 organoid number
200
150
100
50
0
*
tdT Neg
tdT pos
E
tdTomato /DAPI
Muc5b/DAPI
OVX Control
OVX E2
OVX P4
endo
ecto
D
tdTomato /DAPI
Spdef /DAPI
Muc5b /DAPI
Muc1 /DAPI
Non pregnant Mouse Endocervix
Estrus
Metestrus
Diestrus
Epi
100 µm

**Supplemental Figure 4: Olfm4+ progenitors give rise to goblet cells in cervix and vagina in nonpregnant and pregnant states.**

A. RNA expression of tdTomato+ (red), Spdef (cyan) and Muc5b (green) in ectocervical tissue from nonpregnant (diestrus) and GD15. All three markers were detected in the same section, with each signal displayed in separate panels. Images are representative of n≥ 3 biological replicates per time point and scale bars =100 µm.

B. RNA expression of tdTomato+ (red), Spdef (cyan) and Muc5b (green) in vaginal tissue from nonpregnant (diestrus) and GD15. All three markers were detected in the same section, with each signal displayed in separate panels. Images are representative of n≥ 3 biological replicates per time point and scale bars =100 µm.

C. Immunofluorescent staining of tdT positive and tdT negative organoids. Left panels show tdTomato (red) with nuclear counterstain DAPI (blue). Right panels show co-staining for Trp63 (red) and Krt8 (green) with DAPI (blue). Scale bars: 10 µm. Bar graph shows the mean number of organoids formed on Day 12 from tdTomato negative and tdTomato positive cells. Individual points represent n = 3 independent experiments per group. Statistical analysis was performed using a Welch's t-test, with $p<0.05$ considered significant.

D. RNAscope staining of lineage-traced tdTomato$^{+}$ (red) cells with goblet cell markers Spdef (cyan), Muc5b (green), and Muc1 (yellow) across three stages of the nonpregnant endocervix estrus cycle. Representative of n = 3 biological replicates per time point. Scale bars = 100 µm.

E. RNAscope expression of tdTomato$^{+}$ (red) and Muc5b (green) in Ai14Olfm4 lineage OVX control, OVX Estrogen and OVX Progesterone Scale bars = 100 µm.

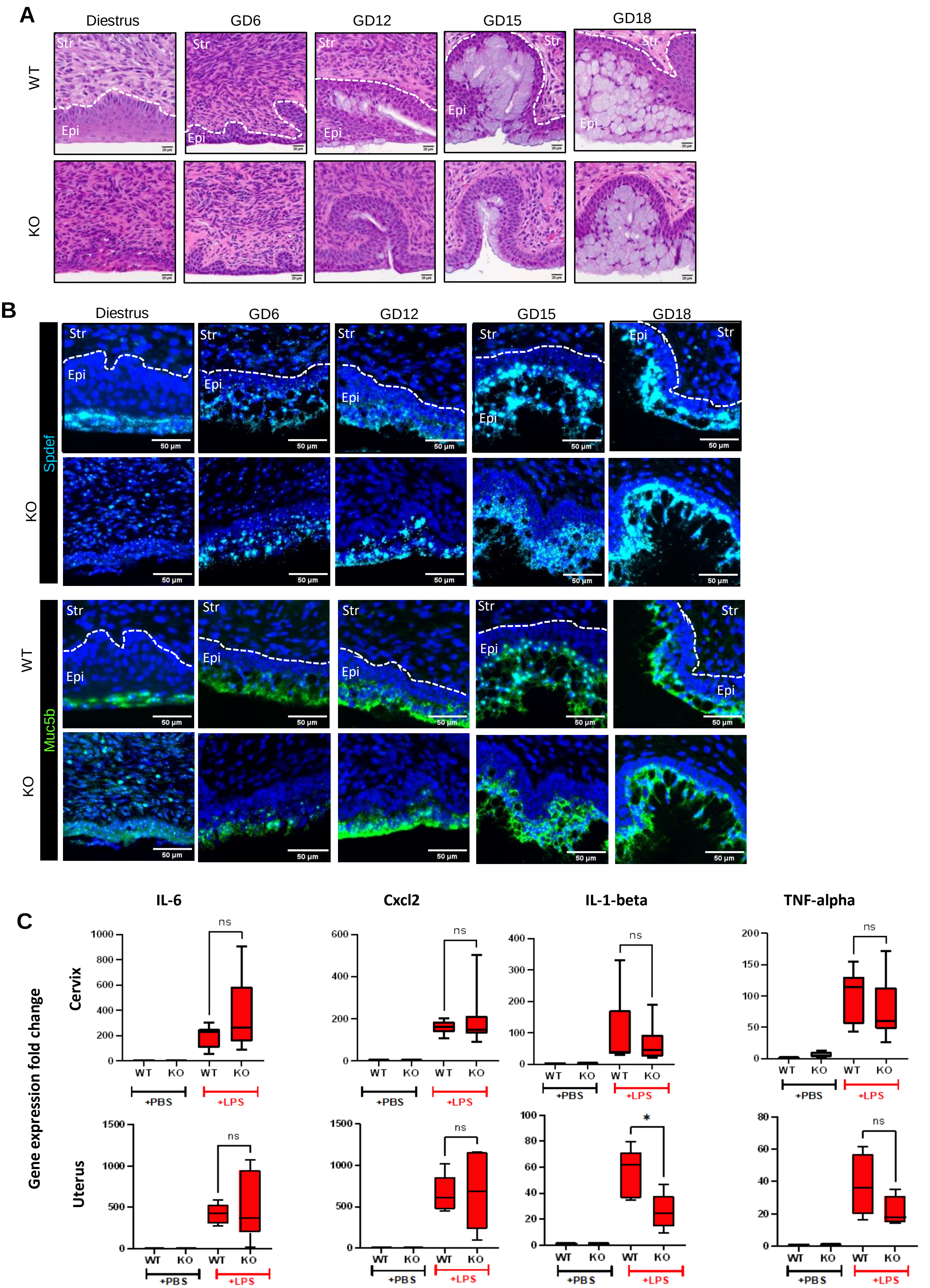
A
Diestrus
GD6
GD12
GD15
GD18
WT
KO
Str
Epi
B
Spdef
Muc5b
50 µm
C
IL-6
Cxcl2
IL-1-beta
TNF-alpha
Cervix
Uterus
Gene expression fold change
WT
KO
+PBS
+LPS
ns
*

**Supplemental Figure 5: Deficiency in Olfm4 does not affect cervical goblet morphology and function**

A. Hematoxylin and eosin (H&E) staining of the endocervical canal in Olfm4 wild-type (WT) and knockout (KO) mice during diestrus and at gestational day 6, 12, 15, 18 timepoints. Representative images from n = 2 biological replicates per genotype and timepoint. Scale bar =20 µm.

B. RNAscope staining for transcription factor Spdef (cyan), Muc5b (green), nuclei (blue) in WT and KO endocervical tissues at nonpregnant diestrus and gestational day 6, 12, 15, 18 timepoints. A single section was stained for the 3 markers and each marker displayed separately in the panel. Representative images from n = 1 biological replicates per condition. Scale bar =50 µm.

C. Gene expression of IL6, Cxcl2, IL1-beta, TNF-alpha inflammatory markers in the cervical (top panel) and uterine tissue (bottom panel) of WT and Olfm4 KO mice exposed to intraperitoneal injection (IP) of lipopolysaccharide (LPS, 5 mg/kg) as compared to IP PBS for 6 hours. (n=5 per condition). Comparisons were performed using the unpaired *t*-test analysis, with $p<0.05$ considered significant.

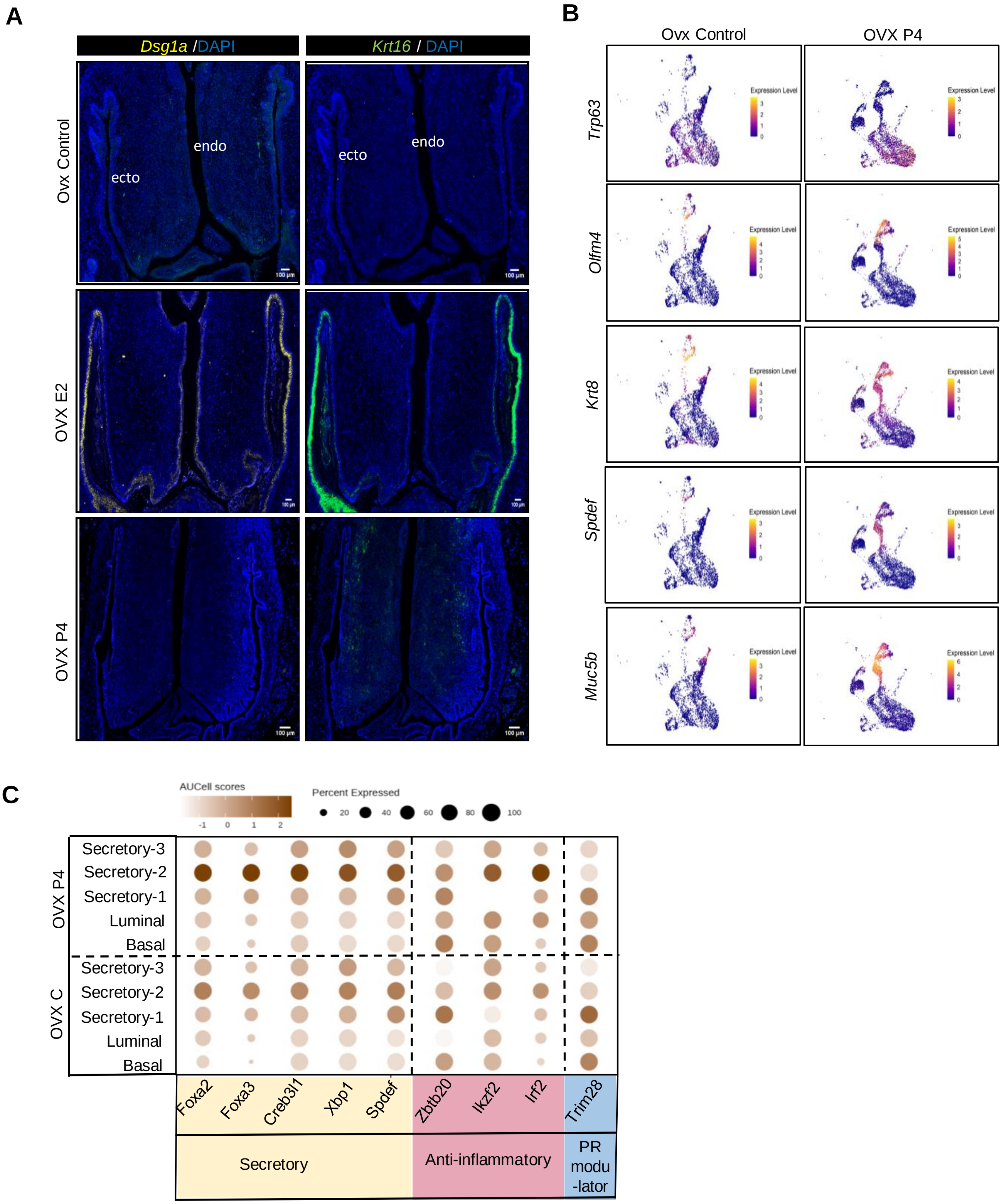
A
Dsg1a /DAPI
Krt16 / DAPI
Ovx Control
OVX E2
OVX P4
ecto
endo
B
Ovx Control
OVX P4
Trp63
Olfm4
Krt8
Spdef
Muc5b
Expression Level
C
AUCell scores
Percent Expressed
20
40
60
80
100
OVX P4
OVX C
Secretory-3
Secretory-2
Secretory-1
Luminal
Basal
Foxa2
Foxa3
Creb3l1
Xbp1
Spdef
Zbtb20
Ikzf2
Irf2
Trim28
Secretory
Anti-inflammatory
PR
modu
-lator

**Supplemental Figure 6: Progesterone induces Olfm4+ progenitor differentiation into goblet cells**

A. RNAscope staining of the lower female reproductive tract (cervix and vagina) for Dsg1a (yellow, left panels) and Krt16 (green, right panels), with nuclei (blue): OVX Control (no hormone), OVX E2 (estradiol), and OVX P4 (progesterone). Both markers were stained on the same sections and are shown in separate panels. Images are representative of n = 3 biological replicates per condition. Scale bars = 100 µm.

B. Feature plots from scRNA-seq showing expression of *Trp63*, *Olfm4*, *Krt8*, *Spdef,* and *Muc5b* across epithelial clusters in OVX Control and OVX P4 cervices.

C. SCENIC analysis predicting transcription factor regulons activated in OVX P4 group epithelial clusters. Transcription factors are grouped by function: secretory associated (cream), anti-inflammatory (pink), modulators of progesterone receptor (PR) activity (blue). The matrix highlights cluster-specific regulon activity, including Foxa2/3, Creb3l1, and Spdef in secretory clusters, Zbtb20 and Irf2 in anti-inflammatory clusters, and Trim28 in PR-modulatory clusters.

| | E. coli ($10^7$ CFU) | | | | IP LPS | |
|---|---|---|---|---|---|---|
| Group | Term | Preterm | PTB rate | n | Mean time to delivery (hours) | n |
| WT | 10 | 10 | 50% | 20 | 13 | 9 |
| KO | 13 | 5 | 28% | 18 | 11 | 9 |

**Supplemental Table 1: Preterm birth outcomes in WT and KO mice following *E.coli* inoculation or LPS treatment. Related to Figure 5.**

Preterm birth outcomes following vaginal inoculation with live pathogenic *E. coli* ($10^7$ CFU) or intraperitoneal injection (IP) of lipopolysaccharide (LPS, 5 mg/kg) on GD15. Fisher's exact test for *E. coli* inoculation: $p = 0.20$; unpaired $t$-test for IP LPS: $p = 0.94$.

| Cell type | Ovx Control | | Ovx P4 | |
|---|---|---|---|---|
| | n Cells | % | n Cells | % |
| Basal | 1354 | 41.0 | 2408 | 37.5 |
| Luminal | 737 | 22.3 | 1174 | 18.3 |
| Secretory-1 | 133 | 4.0 | 1300 | 20.2 |
| Secretory-2 | 298 | 9.0 | 1468 | 22.8 |
| Secretory-3 | 781 | 23.6 | 79 | 1.2 |
| **Total** | **3303** | | **6429** | |

**Supplemental Table 2: Progesterone alters cervical epithelial composition. Related to Figure 6.**

Table summary of epithelial subtype proportions in OVX control vs. OVX P4 samples. The number of cells for each subtype and the percentage of each subtype to the total number of cells is indicated.
